# ER sensing of lipid metabolism drives PRA family-dependent regulation of COPII vesicle transport

**DOI:** 10.1101/2024.11.07.622448

**Authors:** Mitsuki Nakazato, Hiroki Nakamura, Mei Kato, Ryoko Ikema, Mizuki Iguchi, Kazuki Hanaoka, Katsuki Eto, Takefumi Karashima, Asuko Ikeda, Yukari Yabuki, Philipp Schlarmann, Javier Manzano-Lopez, Auxiliadora Aguilera-Romero, Susana Sabido-Bozo, Ana Maria Perez-Linero, Muneyoshi Kanai, Haruyuki Iefuji, Isabelle Riezman, Howard Riezman, Manuel Muñiz, Kouichi Funato

## Abstract

Newly synthesized secretory proteins and many lipids are transported from the endoplasmic reticulum (ER) to the Golgi prior to their ultimate destinations. The ER-to-Golgi transport must be tightly regulated during adaptation to environmental stress. However, the sensing mechanism and regulatory pathways governing the consecutive formation, budding and transportation of COPII vesicles from the ER remain insufficiently explored. Here, we present evidence indicating that COPII-mediated vesicle transport is transcriptionally controlled through the phosphatidic acid-dependent Opi1-Ino2/Ino4 regulatory circuit. Our analysis indicates that *YIP3*, a target gene of Ino2/Ino4, exerts a negative regulatory impact on COPII-mediated vesicle transport. Furthermore, we demonstrated that Ino2/Ino4 but not Yip3 modulates Sar1 activation, the initial step in COPII vesicle formation, whereas Yip3 hinders Sec16 assembly on the ER membrane, thereby implying that Ino2/Ino4 governs COPII-mediated trafficking at multiple steps. Thus, this study provides the first evidence for an ER sensing system that transcriptionally fine-tunes multiple steps of anterograde vesicular transport in response to alterations in lipid composition of the ER membrane.

## INTRODUCTION

Coat protein complex II (COPII) proteins form vesicles that mediate transport of cargo proteins and lipids from the endoplasmic reticulum (ER) to the Golgi apparatus (D’Arcangelo et al., 2013; Venditti et al., 2014; Kurokawa and Nakano, 2019; Bisnett et al., 2021). COPII vesicle formation occurs at a specialized high-curvature domain known as the ER exit site (ERES). ERES is marked by Sec16, which is thought to serve as an essential scaffold for COPII assembly. COPII assembly is triggered by the activation of the small GTPase Sar1 by its guanine nucleotide exchange factor (GEF) Sec12 followed by recruitment of Sec23–Sec24. Subsequently, the Sar1-Sec23-Sec24 complex recruits the Sec13–Sec31 complex that leads to bending of membranes and thereby budding of COPII vesicles from ERES. COPII-mediated trafficking is an essential process in all eukaryotes, and its dysfunction causes a variety of human diseases, including chylomicron retention disease, cranio-lenticulo-sutural dysplasia and several neurological disorders (Venditti et al., 2014; Lu and Kim, 2020; Wang et al., 2020).

Despite its broad physiological significance, important aspects of how COPII-mediated vesicle transport is regulated in response to environmental cues or pathological conditions remain obscure. As efflux of cargo proteins and lipids from the ER depends on demands within the ER, formation of COPII-coated vesicles must be tightly coupled to changes in protein and lipid abundance and composition in the ER membranes. Notably, some lipids contribute to the formation of COPII vesicles by acting directly on membrane deformation, COPII assembly or ERES organization. For example, cone-shaped lipids such as phosphatidic acid (PA), and diacylglycerol (DAG) or inverted cone-shaped lipids such as lysophospholipids are predicted to promote vesicle formation by inducing membrane curvature due to their physical properties (McMahon and Gallop, 2005; Graham and Kozlov, 2010; McMahon and Boucrot, 2015; Harayama and Riezman, 2018). Previous studies have shown that, in mammalian cells, activation of phospholipase D, which catalyzes the formation of PA, is required to support COPII coat assembly and ER export (Pathre et al., 2003). Phospholipase A1 p125, which produces lysophospholipids, has also been shown to be involved in the architecture of ERES (Shimoi et al, 2004; Ong et al., 2010). In mammalian cells, PtdIns4P also acts to support COPII assembly, possibly through the function of PtdIns4P-binding effectors implicated in deforming the membrane (Blumental-Perry et al., 2006). In addition, our previous studies in yeast have found that lysophosphatidylinositol (lyso-PI) increases the recruitment of COPII machinery to membranes and rescue the *sec12-4* mutant by enhancing COPII vesicle formation (Melero et al., 2018). It is also known that loss of phospholipid head groups affects assembly of Sec23 at ERES (Shindiapina and Barlowe, 2010). Thus, curvature-inducing lipids could regulate COPII vesicle formation.

While post-transcriptional modification of COPII coat proteins is utilized as a major mode of regulation of COPII vesicle formation for the immediate response to acute stimuli (Farhan and Rabouille, 2011; D’Arcangelo et al., 2013; Centonze and Farhan, 2019; Bisnett et al., 2021; Kasberg et al., 2023), adaptation of COPII vesicle formation in response to chronic stress due to cargo overload or abnormal organelle homeostasis would be transcriptionally mediated. Thus, an intracellular sensing system is predicted to exist that monitors the membrane state in the ER, transduces signals to the nucleus and controls COPII vesicle formation at the transcriptional level. There is evidence that COPII coat proteins are transcriptionally upregulated in response to ER stress upon physiological nutrient fluctuations or developmental stage in animals (Ishikawa et al., 2017; Farhan et al., 2018; Shaheen, 2018; Liu et al., 2019), suggesting that unfolded protein response (UPR) sensors control COPII vesicle formation via transcriptional regulation of COPII coat proteins. Such transcriptional regulation that transmits from the ER would be conserved for COPII vesicle formation in eukaryotes, because in yeast the *SEC* genes required for COPII vesicle formation are also upregulated by the UPR (Travers et al., 2000). However, the mechanism of transcriptional adaptation for vesicular trafficking coupled with lipid metabolism on the ER membranes is not known.

Here, we show that COPII-mediated vesicle transport is controlled through the Opi1-Ino2/Ino4-dependent transcription regulation, which is modulated by PA levels in the ER. Moreover, our genetic screening combined with gene expression profiling reveals that *YIP3*, a target gene of the transcription factors Ino2/Ino4 is required for the regulation of COPII-mediated trafficking. We also show that Yip3 does not modulate the activation of Sar1, whereas Ino2/Ino4 does, thus suggesting that Ino2/Ino4 regulates COPII-mediated transport at multiple steps. Remarkably, Yip3 hinders Sec16 assembly on the ER membrane. Based on these findings, we propose that this negative regulation serves as a rheostat to adapt COPII vesicle-mediated transport to changes in lipid composition of the ER membrane.

## RESULTS

### Defects in COPII vesicle formation in *sec12-4* mutant are rescued by deletion of *SLC1*, a gene encoding lysophospholipid acyltransferase

The budding yeast *SLC1* gene encodes a lysophospholipid acyltransferase to convert lysophospholipids to phospholipids (Fig. 1 A) (Benghezal et al., 2007; Shui et al., 2010; Henry et al., 2012; Jasieniecka-Gazarkiewicz et al., 2016; Athenstaedt, 2021). Because we showed previously that a *sec12-4* conditional mutation is rescued by lysophospholipid accumulation (Melero et al., 2018), we analyzed if the phenotypes of *sec12-4* temperature-sensitive mutant are suppressed by deletion of *SLC1*. The *sec12-4* mutant strain exhibits growth defects at temperatures above 32°C (Fig. 1 B). Deletion of *SLC1* rescued the growth defects of *sec12-4* mutant cells at restrictive temperatures. Furthermore, testing the maturation of cargo, CPY (carboxypeptidase Y) through the secretory pathway by western blot analysis, we observed that the accumulation of immature forms accumulated in the *sec12-4* mutant was suppressed by *SLC1* deletion (Fig. 1 C, upper panel). As revealed by pulse-chase labeling experiments, a partial rescue of the CPY maturation defect was also seen in the sec*12-4* mutant cells lacking *SLC1* gene (Fig. 1 D). The p2 form of CPY was often invisible. Since it is possible that maturation of CPY may reflect ER-phagy rather than Golgi delivery, we analyzed the maturation of another cargo, Gas1, a glycosylphosphatidylinositol (GPI)-anchored protein. The immature forms of Gas1 accumulated in the *sec12-4* mutant were also suppressed by *SLC1* deletion (Fig. 1 C, lower panel). As *sec12-4* mutant cells have disorganized and huge aggregated ERES and exhibit a reduced number of ERES (Kajiwara et al., 2014; Melero et al., 2018), we next examined whether absence of Slc1 affects the number of ERES in the *sec12-4* mutant. To visualize the ERES, we used the strains expressing *SEC13*-VENUS, a COPII outer coat protein tagged with yellow fluorescent protein (Venus) as an ERES marker (Ikeda et al., 2020). As previously observed (Castillon et al., 2009), the number of Sec13-Venus puncta per *sec12-4* mutant cell was reduced compared to wild-type (Fig. 1 E), and deletion of *SLC1* rescued the phenotype, suggesting that Slc1 negatively regulates COPII vesicle budding from the ER.

**Figure 1.**
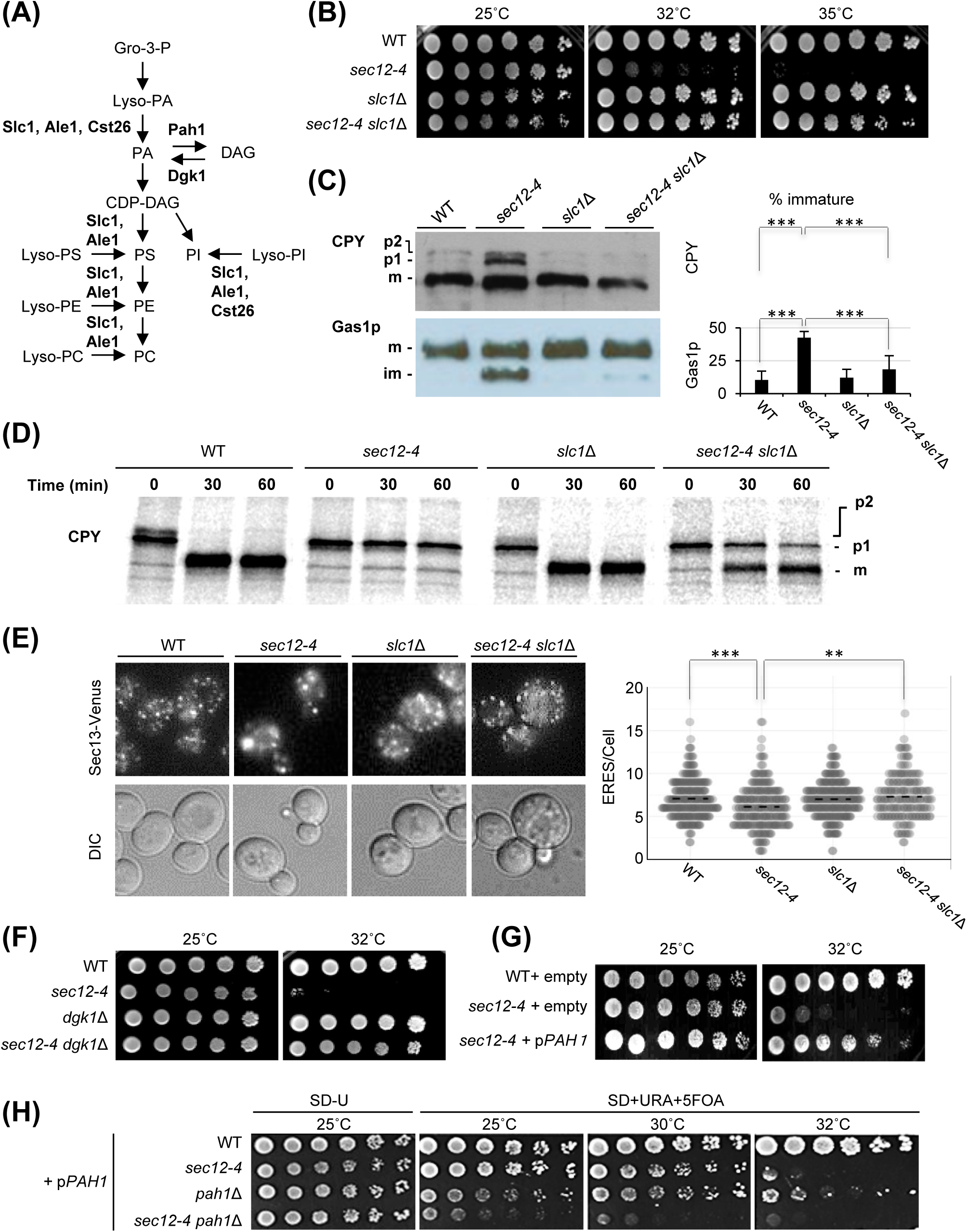
Reduced levels of phosphatidic acid rescue *sec12-4* mutant phenotypes. A) A scheme for the synthesis of phospholipids in yeast. Gro-3-P, glycerol 3-phosphate; Lyso-PA, lysophosphatidic acid; PA, phosphatidic acid; DAG, diacylglycerol; CDP-DAG, cytidine diphosphate diacylglycerol; PI, phosphatidylinositol; PS, phosphatidylserine; PE, phosphatidylethanolamine; PC, phosphatidylcholine; Lyso-PI, lysophosphatidylinositol; Lyso-PS, lysophosphatidylserine; Lyso-PE, lysophosphatidylethanolamine; Lyso-PC, lysophosphatidylcholine. (B) Deletion of *SLC1* rescues temperature-sensitive growth defect of *sec12-4* mutant. Five-fold serial dilutions of cells were spotted onto SD plates and were incubated at the indicated temperatures for 5 days. (C) *slc1*Δ mutation suppresses the accumulation of the ER forms of cargo caused by *sec12-4* mutation. Cell extracts from the cells grown at 32 °C and 25°C were analyzed by western blot for CPY and Gas1, respectively. Mature (m) forms of CPY and Gas1; immature p1 and p2, ER and Golgi forms of CPY, respectively; immature (im) ER form of Gas1 are indicated. Quantification of the percent of immature forms was performed, and the results represent the averages of three independent western blots, and standard deviations are included. Student’s t test: ***p ≤ 0.001. (D) Pulse-chase analysis of CPY was performed by labeling with [^35^S] methionine for 5 min and chasing at 32 °C for the indicated times. CPY was immunoprecipitated followed by SDS-PAGE and analyzed with a phosphoimager. (E) Deletion of *SLC1* suppresses the defect in ERES localization caused by *sec12-4* mutation. Cells expressing Sec13p-Venus were grown at 25°C and observed by fluorescence microscopy. The numbers of fluorescent dots (ERES) per cell were analyzed in three independent experiments, with a minimum of 150 cells counted per genotype. Statistical significance was tested using Tukey-Kramer multiple comparison test: ***p ≤ 0.001, **p ≤ 0.005. (F-H) Five-fold serial dilutions of cells (F), cells transformed with pRS426 (empty) (G) or pRS426-*PAH1* (p*PAH1*) (G, H) were spotted onto SD plates (F), without uracil (G, H) or supplemented with uracil and 5-FOA (H), and were incubated at the indicated temperatures for 5 days. 5-FOA was used to select yeast cells that lose the *URA3* plasmid (p*PAH1*).

This supports a proposed role of lysophospholipids in facilitating COPII vesicle formation (Melero et al., 2018). Since Ale1/Slc4 and Cst26 are also lysophospholipid acyltransferases with different substrate specificity and localization (Chen et al., 2007; Shui et al., 2010; Jasieniecka-Gazarkiewicz et al., 2016; Athenstaedt, 2021), we next tested whether loss of function of their proteins could rescue *sec12-4* mutation. The *ALE1*/*SLC4* disruption suppressed the temperature-sensitive growth defect and CPY maturation defect in the *sec12-4* mutant, whereas the *CST26* disruption did not suppress (Appendix, Fig. S1 A, B and C). These results suggest that Slc1 and Ale1/Slc4 play specific roles in COPII vesicle formation, and are consistent with the findings that Slc1 and Ale1/Slc4 localize to the ER, whereas Cst26 does not localize to the ER but to the lipid particles (Shui et al., 2010; Athenstaedt, 2021).

### Suppression of *sec12-4* mutation by *SLC1* deletion is related to reduced levels of PA

We speculated that deletion of *SLC1* affects the amounts of specific glycerophospholipids, which may participate in the suppression of *sec12-4* phenotypes. Thus, we measured the glycerophospholipid (PC, PE, PI, PS) and lysophospholipid (lyso-PC, lyso-PE, lyso-PI, lyso-PS) composition of cells grown at 32°C with electrospray ionization coupled to triple stage quadrupole (TSQ) vantage mass spectrometry (Dataset EV2 and Appendix, Fig. S2 A and B). We did not analyze PA levels by TSQ vantage mass spectrometer, because other lipids, such as PC could lose their heads in the analysis, leading to false positives for PA. Instead, we analyzed changes in total amount of PA in the cells by thin-layer chromatography (Appendix, Fig. S3 A and B). Neither *sec12-4* or *slc1*Δ mutation changed glycerophospholipid profiles (Appendix, Figs. S2 A, S3 A and B). Interestingly, we detected significant decreases in lyso-PI and lyso-PS in *slc1*Δ or *sec12-4 slc1*Δ strain (Appendix, Fig. S2 B), suggesting that the suppression of *sec12-4* phenotypes by *SLC1* deletion is not related to increased levels of lyso-PI, which facilitate COPII vesicle formation (Melero et al., 2018).

The total amount of PA was not significantly affected by loss of Slc1 (Appendix, Fig. S3 A and B). However, since Slc1 has the capacity to acylate lysoPA to generate PA (Benghezal et al., 2007; Henry et al., 2012; Jasieniecka-Gazarkiewicz et al., 2016; Athenstaedt, 2021), we examined a possible correlation between PA metabolism and the suppression of *sec12-4* mutation by *SLC1* deletion. To test whether a decreased amount of PA could replicate the effect of *SLC1* deletion on temperature-sensitive phenotypes in *sec12-4* mutant, we deleted *DGK1* gene encoding a diacylglycerol (DAG) kinase (Han et al., 2008; Henry et al., 2012), which phosphorylates DAG to produce PA in *sec12-4* mutant cells. At a restrictive temperature 32°C, the *sec12-4 dgk1*Δ cells were able to grow (Fig. 1 F). In addition, we overexpressed *PAH1* that encodes an enzyme catalyzing the dephosphorylation of PA yielding DAG (Han et al., 2006; Henry et al., 2012) in *sec12-4* mutant. Overexpression of *PAH1* rescued *sec12-4* (Fig. 1 G). We also found that *PAH1* deletion aggravates the growth phenotype of the *sec12-4* mutant (Fig. 1 H). These results suggest that reduced levels of PA may contribute to the suppression of *sec12-4* mutation by *SLC1* deletion.

To provide further support for the importance of PA levels, we next studied the localization of Opi1, a negative regulator of UASINO element/ICRE (inositol/choline-responsive element)-dependent transcription, because decreasing PA levels leads to the release of Opi1 from the ER and its entrance into the nucleus (Fig. 2 A) (Loewen et al., 2004; Carman and Henry, 2007; Young et al., 2010; Henry et al., 2012; Fernández-Murray and McMaster, 2016). We took advantage of GFP-tagged Opi1 to address how much PA is present in the ER. Opi1-GFP was largely localized to the perinuclear ER region in the wild-type and *sec12-4* mutant strains, whereas the majority of Opi1-GFP was present in the nucleus of *slc1*Δand *sec12-4 slc1*Δ strains (Fig. 2 B). This result suggests that the level of PA localized to the ER membrane is maintained relatively low in the absence of *SLC1* gene.

**Figure 2.**
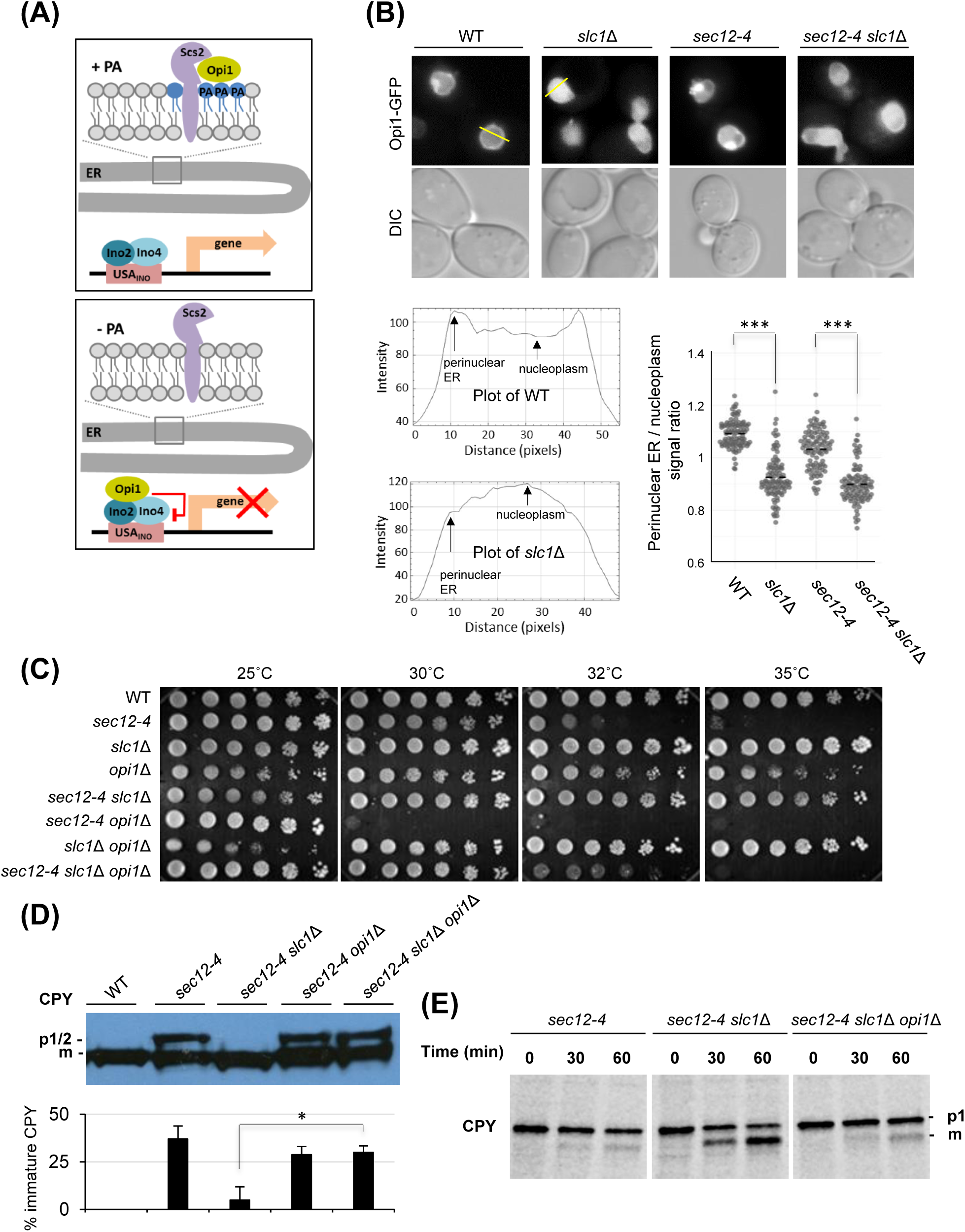
Rescue of *sec12-4* mutant by *SLC1* deletion requires Opi1 transcriptional repressor. (A) A scheme for phosphatidic acid-dependent Ino2/Ino4-Opi1 regulatory pathway. (B) Deletion of *SLC1* leads to the translocation of Opi1-GFP to the nucleus. Cells expressing Opi1-GFP were grown at 25°C and observed by fluorescence microscopy. Bottom panels are intensity plots along the yellow line in wild-type and *slc1*Δ cells. Perinuclear ER/nucleoplasm intensity ratios were analyzed in three independent experiments, with a minimum of 100 cells counted per genotype. Statistical significance was tested using Tukey-Kramer multiple comparison test: ***p ≤ 0.001. (C-E) Spot assay (C), western blot analysis (D) and pulse-chase analysis (E) were performed as described in Figure 1.

### The *OPI1-INO2/INO4* pathway negatively regulates COPII-mediated vesicle transport

Once inside the nucleus, Opi1 binds to Ino2 and attenuates transcriptional activation by the Ino2-Ino4 complex (White et al., 1991; Carman and Henry, 2007; Henry et al., 2012; Fernández-Murray and McMaster, 2016). Therefore, the transcriptional repressor activity of Opi1 may contribute to the suppression of *sec12-4* by *SLC1* deletion. To test this hypothesis, we constructed double (*sec12-4 opi1*Δ) and triple mutant (*sec12-4 slc1*Δ*opi1*Δ), which allowed us to study effects of loss of *OPI1* function on *sec12-4* phenotypes such as temperature sensitivity and CPY maturation. We studied the temperature-sensitive growth phenotype of *sec12-4* mutant. The temperature sensitivity of *sec12-4* was aggravated by *OPI1* deletion, and the suppressive effect of *SLC1* deletion on *sec12-4* growth was also rescued by *OPI1* deletion (Fig. 2 C), suggesting that Opi1 is required for the suppression of *sec12-4* by *SLC1* deletion. Similar results were observed by analyzing the phenotypes in CPY maturation (Fig. 2 D and E).

Because *ALE1*/*SLC4* deletion rescued *sec12-4* mutant phenotypes (Appendix, Fig. S1 A and B), we wanted to determine whether loss of Opi1 function could cancel the effects of *ALE1*/*SLC4* deletion. As expected, the suppressive effect on growth defect of *sec12-4* was rescued by disruption of *OPI1* deletion (Appendix, Fig. S1 D).

We have previously shown that *sec12-4* mutation is rescued by loss of Scs2 and Scs22, the coreceptor for Opi1 (Loewen et al., 2003; Carman and Henry, 2007; Henry et al., 2012; Fernández-Murray and McMaster, 2016), and that the suppressive effect is phenocopied by mutations in Scs2 that are unable to associate with the Opi1 FFAT motif (Kajiwara et al., 2014). Therefore, we tested the effect of *OPI1* deletion on the rescue of growth defects in *sec12-4 scs2*Δ or *sec12-4 scs2*Δ*scs22*Δ. *OPI1* deletion also cancelled the suppressive effect of *scs2*Δ/*scs22*Δ mutation (Appendix, Fig. S1 D). This suggest that the negative regulation of Scs proteins in COPII-mediated vesicle transport is dependent on Opi1.

We next asked if phenotypes of *sec12-4* could be rescued when *INO2* and *INO4* genes were deleted. The temperature-sensitive growth defect of *sec12-4* was restored by *INO2* or *INO4* deletion (Fig. 3 A). Western blot analysis revealed a partial restoration of CPY and Gas1 maturation (Fig. 3 B). Compared to when *SLC1* was deleted (Fig. 1 C), the suppression by *INO4* deletion was less effective. This may reflect a dual role for *SLC1* deletion, one direct on membrane biophysical properties and one indirect via the Ino2/Ino4 pathway. The reduced number of ERES in the *sec12-4* mutant was also rescued by *INO4* deletion (Fig. 3 C). Furthermore, with the same approaches, we examined the effect of *INO4* deletion on temperature-sensitive phenotypes of *sec23-1* and *sec16-2* mutants. Both temperature-sensitive growth defects of *sec23-1* and *sec16-1* mutants were suppressed in the absence of Ino4 (Fig. 3 D). Cargo maturation defects in *sec23-1* were also restored (Fig. 3 E). Taken together, these results suggest that *OPI1*-*INO2*/*INO4* pathway negatively regulates COPII-mediated trafficking.

**Figure 3.**
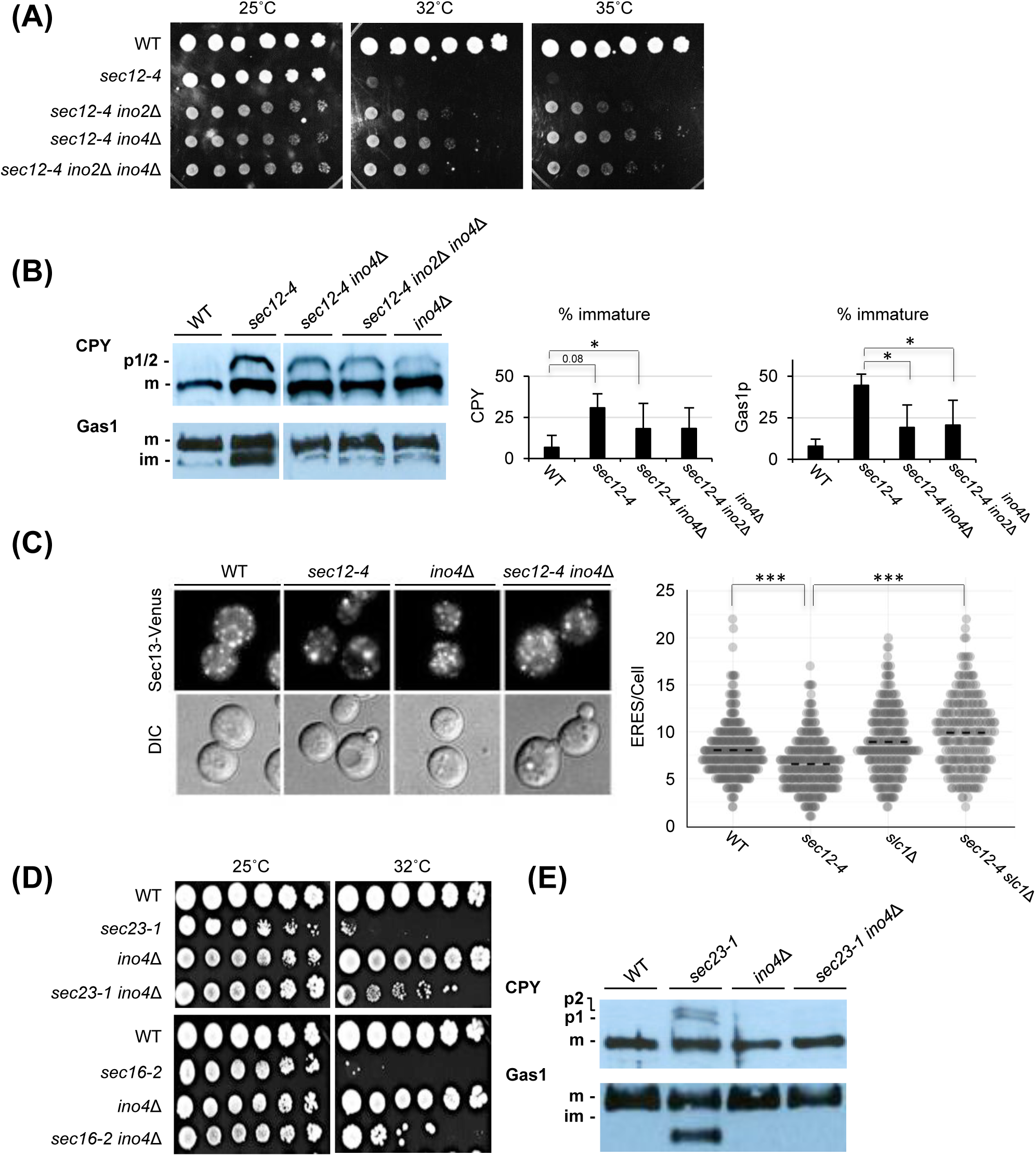
Deletion of *INO2* and *INO4* rescues sec mutant phenotypes. (A-E) Spot assay (A, D), western blot analysis (B, E) and fluorescence analysis of ERES localization at 32°C with a minimum of 250 cells counted per genotype (C) were performed as described in Figure 1. Student’s t test (B): *p ≤ 0.05 and Tukey-Kramer multiple comparison test (C): ***p ≤ 0.001.

### Screening for candidate genes involved in regulating COPII-mediated vesicle transport

To identify genes that negatively regulate COPII-mediated vesicle transport via Ino2/Ino4-dependent transcription, we analyzed the transcriptomic profile of *sec12-4 ino2*Δ *ino4*Δ mutant and compared it to the profile of *sec12-4* mutant cells (Dataset EV3). Expression of 152 known genes was significantly downregulated by more than twofold upon deletion of *INO2*/*INO4* and the functional categorization showed that different functional categories of genes were regulated by the *OPI1*-*INO2*/*INO4* pathway (Fig. 4 A). Consistent with previous reports (Santiago and Mamoun, 2003; Jesch et al., 2005), the expression of *OPI3* (relative amounts are 0.52±0.01, p<0.001, mean ± s.d, n=3), *CHO1* (0.56±0.13, p<0.05) and *CHO2* (0.67±0.09, p<0.05) that contain the UASINO element in their promoters was significantly down-regulated compared to the expression level of control strain, although the decreases in expression were not more than 2-fold. In addition, we found that there are no significant changes in expression of genes involved in phospholipid metabolism such as *SLC1*, *ALE1*, *DGK1*, *PAH1*, *PLB1*, *PLB2* and *PLB3*, except *NTE1*, and genes functioning upstream of *INO2*/*INO4* such as *SCS2*, *SCS22* and *OPI1* (Appendix, Fig. S4 A). These results suggest that suppression of *sec12-4* mutation by *INO2*/*INO4* deletion is not due to changes in their gene expression. We also did not observe significant 1.5-fold or greater increases in expression of *SEC* genes involved in COPII vesicle formation (Appendix, Fig. S4 B), implying that other genes are involved in the *OPI1*-*INO2*/*INO4* pathway-dependent regulation of COPII vesicle formation.

**Figure 4.**
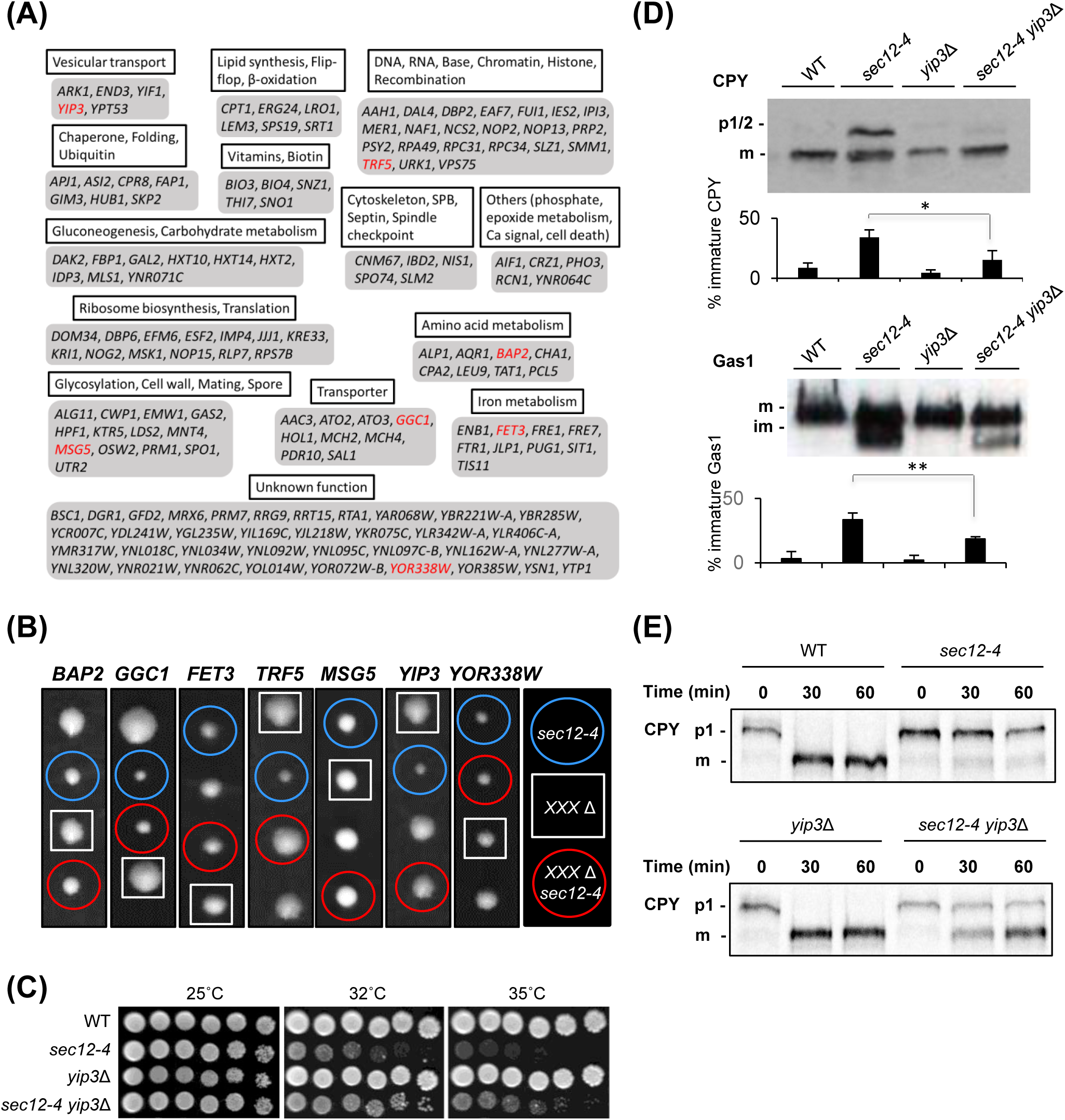
Deletion of *YIP3*, a target gene of Ino2/Ino4, suppresses *sec12-4* mutant phenotypes. (A) Functional groups among 152 genes whose expression increased more than twofold in *sec12-4 ino2*Δ *ino4*Δ cells compared to *sec12-4* cells. 7 genes that cause a growth defect when overexpressed in wild-type cells (Appendix, Figs. S5 and S6) are shown in red. (B) Tetrad analysis reveals that deletion of *TRF5* or *YIP3* suppresses the slow growth defect of *sec12-4* mutant. Diploids heterozygous for both *sec12-4* and disruption of 7 candidate genes for negative regulators of COPII vesicle formation shown in red (A) were sporulated, and the tetrads were dissected and incubated on YPD plates for 5 days. Blue circles, squares and red circles represent *sec12-4*, *XXX*Δ and *sec12-4 XXX*Δ haploid cells, respectively. (C-E) Spot assay (C), western blot analysis (D) and pulse-chase analysis at 30 °C (E) as described in Figure 1 shows rescues of *sec12-4* mutant by *YIP3* deletion. Student’s t test (D): **p ≤ 0.01, *p ≤ 0.05.

Thus, we screened for the 152 downregulated genes to identify the genes that cause growth defects when overexpressed (Appendix, Fig. S5), because the genes of interest negatively regulate the formation of COPII vesicles. For this purpose, we used a gene expression method, named genetic tug-of-war (gTOW) that causes the overexpression of the target gene due to an increase in the copy number, which is inversely correlated with the leucine concentration in the growth medium (Moriya et al., 2006). We found that overexpression of 7 genes, including *BAP2*, *GGC1*, *FET3*, *TRF5*, *MSG5*, *YIP3*, and *YOR338W*, led to growth defects in wild-type cells of BY background strain (Fig. 4 A, shown in red and Appendix, Figs. S5 and S6). To validate and refine the candidate genes obtained by our primary screen, we next investigated whether disruption of those genes could rescue the growth defect of *sec12-4* mutant, because *sec12-4* cells grow slowly in comparison to wild-type cells at permissive temperature (25°C). We generated heterozygous diploid double mutant BY background strains, sporulated and subjected to tetrad dissection. After spores were allowed to form colonies for several days, we determined the colony size and genotypes. The colonies of *sec12-4 trf5*Δ and *sec12-4 yip3*Δ double mutants were larger than that of single *sec12-4* mutant (Fig. 4 B), indicating that deletion of *TRF5* or *YIP3* rescues the slow growth of *sec12-4* mutant. Furthermore, we observed that deletion of those genes partially suppressed the temperature-sensitive growth defects of *sec12-4* mutant (Appendix, Fig. S7). Thus, we identified two potential genes that are responsible for regulation of COPII-mediated vesicle transport via the *OPI1*-*INO2*/*INO4* pathway.

### *YIP3* upregulated by Ino2/Ino4 negatively regulates COPII-mediated vesicle transport

Trf5 is a noncanonical poly(A) polymerase and assembles into the heteromeric Trf5-Air1-Mtr4 (TRAMP) complex that promotes the degradation of RNA substrates by interacting with the nuclear exosome (LaCava et al., 2005; Schmidt and Butler, 2013; Bresson and Tollervey, 2018). Because *TRF5* deletion increases RNA abundance for RNAs showing Trf5 binding (Delan-Forino et al., 2020), the most straightforward explanation, which fits with our findings, could lie in the role of TRAMP complex in RNA levels. Hence, increased expression levels of proteins involved in the formation of COPII vesicles might be a reason for the suppression of *sec12-4* mutation by deletion of *TRF5*.

On the other hand, *YIP3* encodes a protein known to localize to COPII vesicles (Otte et al., 2001) and has been proposed to be involved in vesicular transport between the ER and Golgi apparatus because it physically interacts with yeast Rab proteins (Calero and Collins, 2002; Geng, et al., 2005), but its function is unknown. No changes in the localizations of yeast Rab proteins and in the maturation of CPY were observed in a *yip3*Δ mutant, whereas overproduction of Yip3 caused a severe growth defect (Appendix, Figs. S5 and S6) and aberrant ER membrane proliferation (Geng, et al., 2005), suggesting the role of Yip3 as a negative regulator of ER homeostasis. As ER membranes have been shown to proliferate in mutants defective in COPII vesicle formation (Kaiser and Schekman, 1990), we therefore examined whether Yip3 negatively regulates COPII-mediated vesicle transport. First, we created a *sec12-4 yip3*Δ double mutant with a different genetic background than BY and confirmed that the suppressive effect of *YIP3* deletion on growth defect of *sec12-4* is not background specific (Fig. 4 C). Next, we tested whether immature cargos accumulated in the *sec12-4* mutant could be restored by *YIP3* deletion. Western blot analysis revealed the restoration of CPY and Gas1 maturation by *YIP3* deletion (Fig. 4 D). Furthermore, pulse-chase labeling experiments showed that CPY maturation defect in *sec12-4* mutant is partially restored in the absence of Yip3 (Fig. 4 E). These results suggest that Yip3 negatively regulates COPII-mediated vesicle transport. Thus, when the results are taken together, we conclude that the *OPI1*-*INO2*/*INO4* pathway in response to changes in PA metabolism regulates anterograde vesicular transport from the ER via *YIP3* transcription.

### The PA-Opi1-Ino2/Ino4 system plays a critical role in controlling COPII vesicle formation by tuning multiple steps

What roles can Yip3 play in COPII-mediated trafficking? Because phenotypes of *sec12-4* mutant, which is defective in GDP-to-GTP exchange on Sar1, have been shown to be rescued by Sar1 overexpression (Nakańo A and Muramatsu, 1989; d’Enfert et al., 1991; Van der Verren and Zanetti, 2023), it is possible that the suppression by *YIP3* deletion is due to an increased level of Sar1 protein, although no effect of *INO2*/*INO4* disruption on mRNA levels of *SAR1* gene was observed (Appendix, Fig. S4 B). Therefore, we first studied the protein levels of Sar1. Western blot analysis showed that the steady-state levels of Sar1 were unaffected by *YIP3* deletion (Appendix, Fig. S8 A) as well as by *INO4* deletion (Appendix, Fig. S8 B).

We next tried to determine if Yip3 functions to inhibit the GDP-to-GTP exchange on Sar1. Since Sar1 associates with the ER membrane upon activation from the GDP- to the GTP-bound form of by Sec12 (Nakańo A and Muramatsu, 1989; d’Enfert et al., 1991; Van der Verren and Zanetti, 2023), we assessed the level of association of Sar1-AcGFP, a C-terminally AcGFP-fused wild-type Sar1 that cycles between the GDP/GTP-bound states to the ER membrane in vivo (Yorimitsu and Sato, 2012). In wild-type cells, ER membrane/cytosol signal ratios (an index of ER membrane association) of Sar1-AcGFP were between 1.3 and 1.5 (Fig. 5 A). On the other hand, as expected, the ratios of Sar1D32G-AcGFP, a GDP- locked form of Sar1, were significantly reduced to about 1.1. A reduction in the ER membrane association of Sar1-AcGFP was also observed in the *sec12-4* mutant strain. These results are consistent with the model that activated Sar1-GTP associates with the ER membrane. We analyzed the ER membrane association of Sar1-AcGFP in the *sec12-4 yip3*Δ mutant and showed that *YIP3* deletion fails to rescue the ER membrane association defect in *sec12-4* mutant (Fig. 5 B), suggesting that Yip3 is not involved in regulating Sar1 activity. Interestingly, we found that the ER membrane association defect in *sec12-4* mutant was partially rescued by loss of Ino2 or Ino4 (Fig. 5 C). This result suggests that there is an unknown factor in the Ino2/Ino4 target genes that functions to negatively regulate the ER membrane association of Sar1. Taken together, these results support a critical role of the PA-Opi1-Ino2/Ino4 system in regulating COPII vesicle formation by tuning multiple steps.

**Figure 5.**
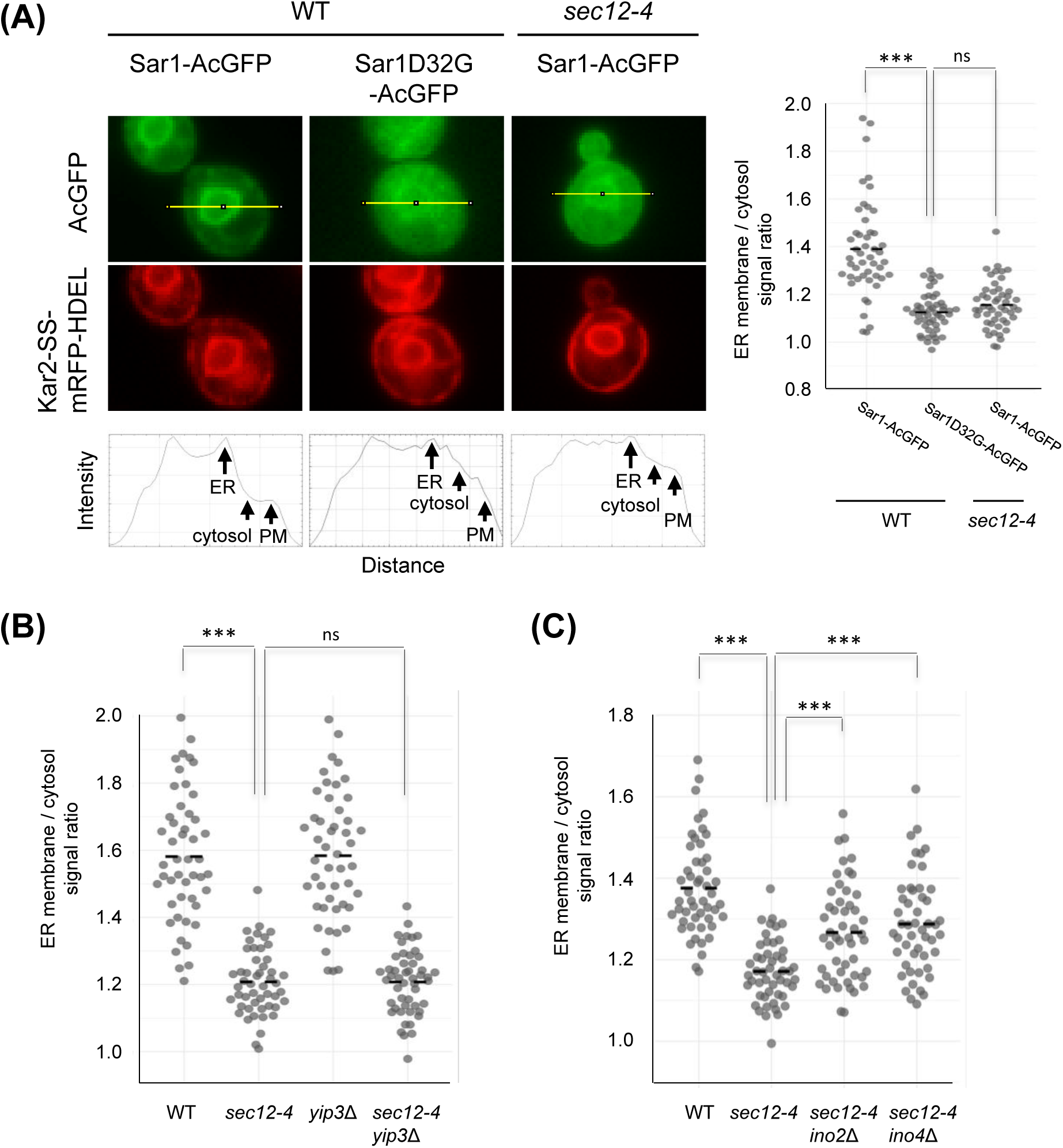
Deletion of *INO2* or *INO4* but not *YIP3* rescues Sar1 membrane association defect in *sec12-4* mutant. (A-C) Cells expressing Sar1-AcGFP or Sar1D32G-AcGFP and Kar2-SS-mRFP-HDEL were grown at 25°C in SD medium, shifted to 32 °C for 60 min and observed by fluorescence microscopy (left panel of A). Bottom panels are intensity plots along the yellow line in wild-type and *sec12-4* cells. ER membrane/cytosol intensity ratios were analyzed in three independent experiments, with a minimum of 50 cells counted per genotype (right panel of A, B and C). Statistical significance was tested using Tukey-Kramer multiple comparison test: ***p ≤ 0.001, ns not significant.

### Yip3 is involved in the regulation of Sec16 assembly on the ER membrane

Yip3 was not involved in the regulation of Sar1 activity. Therefore, it was considered that Yip3 may function in the later stages of the Sar1 activity. Sar1 on the ER membrane recruits the inner coat Sec23/Sec24 complex from the cytosol, which than recruits the outer coat Sec13–Sec31 complex to deform the membrane and form COPII vesicles (D’Arcangelo et al., 2013; Venditti et al., 2014; Kurokawa and Nakano, 2019; Bisnett et al., 2021). As Sec16 interacts with all of the COPII coat subunits, it is thought to act as a scaffold for COPII assembly to form ERES (Barlowe and Miller, 2013). Sec16 is also proposed to act as a Sec23 GAP inhibitor to contribute to stable coat assembly. A recent study has shown that Sec16 localizes to the ERES in a manner that is dependent on Sed4 (Yorimitsu and Sato, 2023). To characterize whether Yip3 is associated with Sec16 function, we tested colocalization for Sec16 with Yip3. For this purpose, we generated a wild-type strain that express Sec16-GFP and Yip3-mCherry from their endogenous loci and visualized by fluorescence microscopy. Like Sec16-GFP, consistent with a previous report with Yip3-GFP (Geng et al., 2005), Yip3-mCherry showed localization in puncta (Fig. 6 A). Approximately 5 % and 20 % of puncta containing Sec16-GFP were overlapped and closely associated with Yip3-mChrrey puncta, respectively, suggesting that a portion Yip3 colocalizes with Sec16 on the ER membrane.

**Figure 6.**
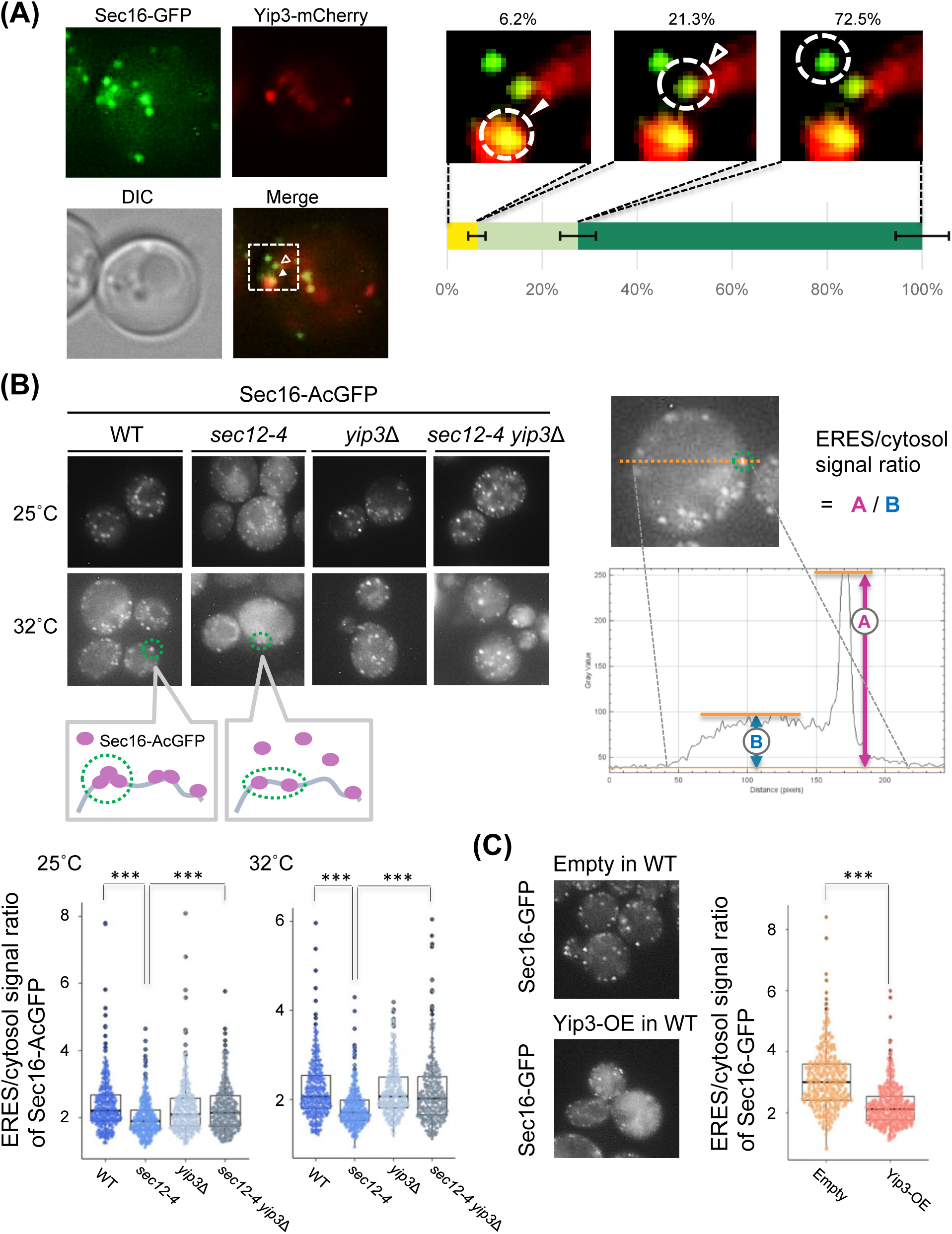
Yip3 inhibits Sec16 assembly into the ERES. (A) Wild-type cells expressing endogenously tagged Sec16-GFP and Yip3-mCherry were grown at 25°C in YPD medium, and observed by fluorescence microscopy. Sec16-GFP puncta were classified as “overlapped”, “closely associated” with Yip3-mCherry puncta, or “away” from Yip3-mCherry puncta. White closed and opened arrowheads indicate areas of Sec16-GFP overlapped and closely associated with Yip3-mCherry, respectively. At least 1000 of Sec16-GFP puncta were evaluated per experiment, and the results represent the averages of three independent experiments, and standard deviations are included. (B) Cells expressing Sec16-AcGFP were grown at 25°C in SD medium, shifted to 32 °C for 60 min and observed by fluorescence microscopy. The fluorescence intensities of Sec16-AcGFP in ERES (puncta, A) and in cytosol (B) were measured (Yorimitsu and Sato, 2023). After subtracting the background values, the ratio of intensities in ERES and cytosol (A / B) was analyzed in three independent experiments, with a minimum of 300 of Sec16-AcGFP puncta counted per genotype. Statistical significance was tested using Tukey-Kramer multiple comparison test: ***p ≤ 0.001. (C) Yip3 was overexpressed in wild-type cells expressing endogenously tagged Sec16-GFP. The empty vector was used as a control. After grown at 25°C in SD medium, cells were observed by fluorescence microscopy. The ratio of intensities of Sec16-GFP in ERES and cytosol was determined as described in (B). Tukey-Kramer multiple comparison test: ***p ≤ 0.001.

To more directly determine whether Yip3 is involved in Sec16 function, we assessed the assembly of Sec16 to the ERES (Yorimitsu and Sato, 2023). Interestingly, we found that in wild-type cells, Sec16-AcGFP displayed little signal in the cytosol, whereas it was detected in the ERES but also showed a diffuse cytoplasmic signal in *sec12-4* mutant cells (Fig. 6 B). We reasoned that if Yip3 hinders Sec12-dependent assembly of Sec16 into the ERES, deletion of *YIP3* would rescue the Sec16 localization defect in the *sec12-4* mutant. Indeed, we observed that the Sec16 localization defect was fully restored by *YIP3* deletion as evidenced by increased ERES/cytosol signal ratio of Sec16-AcGFP, suggesting that Yip3 has the ability to inhibit the assembly of Sec16 into the ERES. To verify this, we assessed the impact of Yip3 overexpression on Sec16 assembly in the wild-type cells. We confirmed that overexpressing Yip3 hinders Sec16 assembly into the ERES (Fig. 6 C). Collectively, our findings suggest that Yip3 saves as a negative regulator of COPII vesicle formation that inhibits Sec16 assembly into the ERES.

## DISCUSSION

Although the biochemical properties of the formation of COPII vesicles and the fusion with their target membranes are relatively well known (D’Arcangelo et al., 2013, Venditti et al., 2014; Kurokawa and Nakano, 2019; Bisnett et al., 2021), less is known about how the COPII vesicle-dependent transport system adapts to changes in physiological cues and stress. Post-transcriptional modifications have emerged as major modes of regulation of the COPII pathway (Farhan and Rabouille, 2011; D’Arcangelo et al., 2013; Centonze and Farhan, 2019; Bisnett et al., 2021; Kasberg et al., 2023). Nevertheless, extensive regulation occurring at the transcriptional level must also be important for environmental adaptation. In this study, we provide evidence that PA sensing in the ER transcriptionally regulates COPII-mediated vesicle transport. Our data suggest that a decreased PA in the ER membranes suppresses the secretory defect of the *sec12-4* mutant. This suppression is caused by inhibiting the Ino2/Ino4 transcriptional activator complex through its interaction with Opi1, which is sequestered in the ER by PA and when released from the ER, is translocated into the nucleus to repress Ino2/Ino4 target genes. Furthermore, we found that the transcriptional regulation of COPII-mediated vesicle trafficking requires *YIP3*, an Ino2/Ino4 target gene. We show that Yip3 deletion restores the secretory defect of *sec12* mutant, suggesting that Yip3 negatively regulates COPII-mediated vesicle transport. This readily explains the negative effects of Yip3 overexpression on yeast cell growth (Appendix, Figs. S5 and S6) and ER morphology (Geng et al., 2005). Consistent with the role of Yip3 as a negative regulator, we found that Yip3 inhibits Sec16 assembly into the ERES (Fig. 6 C). Thus, our findings suggest that Yip3 negatively regulates COPII vesicle formation via transcriptional expression by the PA-Opi1-Ino2/Ino4 pathway (Fig. 7). We propose that the PA-Opi1-Ino2/Ino4-Yip3 pathway may serve as a rheostat linking the ER membrane status to COPII vesicle formation.

**Figure 7.**
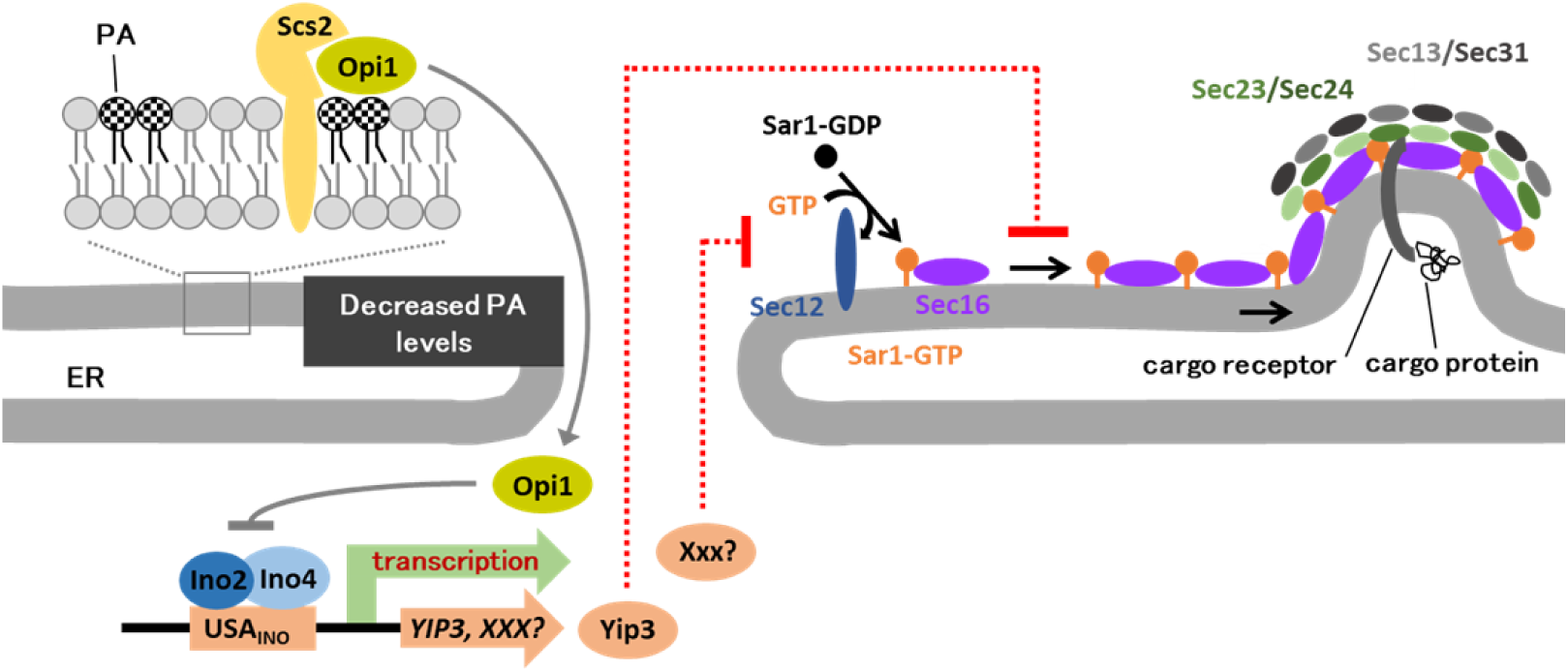
A model of ER sensing of phosphatidic acid (PA) metabolism in regulating COPII vesicle formation. Yip3 and Xxx (unknown protein) encoded by *YIP3* and *XXX*, target genes of the Ino2/Ino4, act as negative regulators of COPII vesicle formation. When PA levels in the ER membranes decrease, Opi1 is translocated to the nucleus, binds to Ino2 and attenuates transcriptional activation by the Ino2/Ino4 complex. This leads to the repression of Yip3 and Xxx expression and hence the release of negative regulation of COPII vesicle formation. The fact that deletion of *INO2* or *INO4* but not *YIP3* rescues Sar1 membrane association defect in *sec12-4* mutant reveals that Yip3 and Xxx proteins regulate different steps in COPII vesicle formation. Yip3 hinders Sec16 assembly into the ERES.

Loss of *SLC1* did not affect the total amount of phospholipid species (Appendix, Figs. S2 A, S3 A and B). Given the roles of Slc1 in the fatty acid remodeling of phospholipids (Benghezal et al., 2007; Shui et al., 2010; Henry et al., 2012; Jasieniecka-Gazarkiewicz et al., 2016; Athenstaedt, 2021), we asked how the lack of *SLC1* influences fatty acyl chain composition of phospholipids. As previously reported (Shui et al., 2010), relative amounts of phospholipid species such as 26:0, 26:1, 26:2, 28:0, 28:1 and 28:2 composed of short fatty acids decreased (Dataset EV4 and Appendix, Fig. S9). This suggests that Slc1 preferentially utilizes short fatty acids as substrates to remodel phospholipids. Remarkably, *slc1*Δ strain showed decreased levels of lyso-PI and lyso-PS (Appendix, Fig. S2 B). Decreases in lyso-PI 16:0 and lyso-PI 18:1 species, the majority of lyso-PI, have also been observed in *slc1*Δ cells (Shui et al., 2010). Because Slc1 has preferential substrate specificity for PI and PS, like PA (Benghezal et al., 2007), the decreases in lyso-PI and lyso-PS in *slc1*Δ cells may be related to the substrate specificity of Slc1. Although local abundance of lysophospholipids in the ER membrane is unknown, the finding that *SLC1* deletion does not increase the total amounts of lysophospholipids such as lyso-PI, -PS, -PE and -PC suggests that suppression of the *sec12-4* mutant is not due to their increased lysophospholipids, which facilitate COPII vesicle formation as proposed previously (Melero et al., 2018). Like lysophospholipids, PA has an unique physical property that generates membrane curvature (McMahon and Gallop, 2005; McMahon and Boucrot, 2015; Harayama and Riezman, 2018), thereby facilitating the budding of vesicles. To determine the local abundance of PA in the ER, we analyzed the binding of Opi1-GFP to the ER membrane by fluorescence microscopy, and found that the amount of Opi1-GFP bound to the membrane decreased in the absence of Slc1. This suggests that deletion of *SLC1* reduces local levels of PA in the ER, giving rise to the possibility that a local increase in lyso-PA in the ER may be responsible for the suppression of the *sec12-4* mutation. However, the suppression by *SLC1* deletion was cancelled by loss of Opi1, implying that the negative effect of Slc1 on COPII-mediated vesicle transport is dependent on Opi1 rather than local increase in lyso-PA level.

Decreased PA dissociates Opi1 from the ER and thereby represses expression of Ino2/Ino4 target genes through binding of Opi1 to the Ino2 (Carman and Henry, 2007; Henry et al., 2012; Fernández-Murray and McMaster, 2016). Consistently, we showed that deletion of *INO2*/*INO4* also rescued the *sec12-4* mutation, suggesting that COPII-mediated vesicle transport is negatively regulated by genes that are transcriptionally upregulated by Ino2/Ino4. The PA-Opi1-Ino2/Ino4 system needs to maintain ER homeostasis during ER stress (Schuck et al., 2009; Papagiannidis et al., 2021). ER stress upregulates both genes involved in protein folding and degradation and genes required for COPII vesicle formation (Travers et al., 2000). This suggests that UPR acts to reduce the level of misfolded protein not only by promoting folding to the natural state and increasing the rate of protein degradation, but also by facilitating their exit from the ER via the secretory pathway, presumably to be degraded in the vacuole. The latter is consistent with the evidence that mutations affecting COPII vesicle formation have negative genetic interactions with mutations conferring defects in UPR (Chang et al., 2004) and that the constitutive activation of the UPR could suppress *sec12-4* mutant phenotypes (Sato et al., 2002). In mammals, it has also been shown that the IRE1 and XBP1, the key regulators of the UPR, are functionally connected to COPII-mediated trafficking, and transcriptional regulation of COPII components by XBP1 contributes to the promotion of COPII-dependent trafficking (Liu et al., 2019). However, our DNA microarray analysis in the *sec12-4* mutant background showed that disruption of *INO2*/*INO4* hardly changed the expression of UPR target genes encoding a chaperone (*SCJ1*; 1.09±0.07 fold, p=0.16) and required for ERAD (*DER1*; 1.15±0.06 fold, p=0.05). Moreover, we observed that *SEC* genes required for COPII vesicle formation are not significantly upregulated in the absence of Ino2/Ino4. These results suggest that the suppression of *sec12-4* mutation by Ino2/Ino4 deletion is independent of the activity of the UPR signaling pathway and is not due to increased expression of COPII components. By genetic screening combined with gene expression profiling, we identified *YIP3* as a negative regulator of COPII-dependent trafficking. *YIP3* was downregulated more than 2-fold in the absence of Ino2/Ino4. This is in agreement with a previous report showing that *YIP3* is an Opi1-regulated gene (Jesch et al., 2005). *YIP3*-encoded protein Yip3 is localized to COPII vesicle fraction, the ER and the Golgi, and are thought to function in transport through the early secretory pathway, but the role of Yip3 was unknown (Otte et al., 2001; Calero and Collins, 2002; Geng et al., 2005; Shaik et al., 2019; Shaik et al., 2019; Angelotti et al., 2022). Our experiments demonstrated that *YIP3* deletion rescues *sec12-4* mutant phenotypes, suggesting that Yip3 negatively regulates COPII-mediated vesicle transport and transcriptional repression of *YIP3* is the main mechanism of suppressing the *sec12-4* phenotypes in the absence of Ino2/Ino4.

What is the function of Yip3? Yip3 is the only yeast protein belonging to the PRA family, which is named for its role as a ‘Prenylated Rab acceptor’, and is highly conserved from yeast to humans (Angelotti et al., 2022). Human PRA1 was proposed to function as a GDI-displacement factor that dissociates Rab GTPases from Rab-GDP-dissociation inhibitor GDI (Sivars et al., 2003). Like PRA1, Yip3 interact with a member of the yeast Rab family Ypt1 (Calero and Collins, 2002; Geng et al., 2005), which is involved in ER-to-Golgi trafficking. However, distribution of Ypt1 between the cytosol and membranes was not affected in Yip3-deleting or -overproducing cells (Geng et al., 2005), suggesting that the negative regulation of Yip3 in ER-to-Golgi trafficking found in this study may not be linked to a mechanism that requires Ypt1. This is not inconsistent with evidence that Ypt1 is not required for COPII vesicle formation (Rexach and Schekman, 1991; Heidtman et al., 2003). If, like Yip3, the other PRAF family negatively regulates COPII vesicle-mediated transport from the ER, its expression would result in ER accumulation of immature cargo proteins. Indeed, immature glycosylation patterns of the glutamate transporter have been observed in mammalian cells expressing PRA family member, PRA2/GTRAP3-18 (Ruggiero et al., 2008). Similar immature glycosylation of cargo proteins was observed in Arabidopsis expressing the PRA isoform, AtPRA1.B6 (Lee et al., 2011). While *YIP3* deletion failed to rescue the ER membrane association defect of Sar1 in *sec12-4* mutant, it rescued the Sec16 localization defect in *sec12-4*. These results suggest that Yip3 acts to inhibit Sar1-dependent Sec16 assembly at the ERES (Fig. 7). This was confirmed by the results with Yip3 overexpression. Furthermore, this proposal is consistent with a previous report that Yip3 overexpression causes ER membrane expansion and growth arrest (Geng et al., 2005). Recently, it was demonstrated that Sed4, a homolog of Sec12, is essential for the assembly of Sec16 to ERES through its interaction with Sec16 (Yorimitsu and Sato, 2023). Thus, Yip3 might impede the assembly of Sec16 to ERES by inhibiting the binding between Sed4 and Sec16. Otherwise, since Yip3 physically interacts with Sec23, Sec24 and Sec13 (Graef et al., 2013), it is possible that Yip3 may disturb Sec16 localization to ERES through its interaction with the COPII components. It would be interesting to know whether mammalian homologs of the Yip3/PRA family affect Sec16 localization.

Another striking feature from this study is that the defect in ER membrane association of Sar1 in *sec12-4* mutant was rescued by loss of Ino2 or Ino4. This suggests that target genes of Ino2/Ino4 contain a factor that regulates Sar1 activity. Because loss of Ino2/Ino4 did not increase the mRNA levels of *SEC12* and because *YIP3* deletion failed to rescue the defect in the ER membrane association of Sar1 in *sec12-4* mutant, an unknown factor distinct from them may be involved in the activation of Sar1 (Fig. 7). Alternatively, it is possible that structural changes in the ER membrane by loss of Ino2 or Ino4 affect Sar1 activity, since Ino2/Ino4 complex is required for ER membrane expansion (Schuck et al., 2009). Thus, we propose that the PA-Opi1-Ino2/Ino4 system fine tunes COPII vesicle formation by coordinating multiple steps in response to changes in the lipid composition of the ER membrane.

Finally, our study highlights the identification of an unexpected player, Yip3 as a negative regulator of COPII-mediated vesicle transport. Moreover, the findings reported herein reveal that lipid sensing in the ER controls COPII-mediated vesicle transport via transcriptional repression of Yip3. Because Yip3 is highly conserved from budding yeast to human (Angelotti, 2022), it is likely that such a mechanism for Yip3-mediated lipid-based transcriptional regulation may exist in higher organisms. Although understanding the regulatory mechanism of COPII-mediated trafficking by Yip3 remains as an important challenge for the future and an unknown factor that is the target of Ino2/Ino4 and negatively regulates Sar1 activity remains to be identified, this study opens up new avenues of research in elucidating the regulatory mechanisms of COPII vesicle formation.

## MATERIALS AND METHODS

### Yeast strains and plasmids

All strains of S. cerevisiae used in this study are listed in Dataset EV1. Double mutants were constructed by crossing haploid yeast strains containing single-gene mutations in same backgrounds, sporulation and subsequent dissection of the spores. The genotypes of spores were verified by colony PCR. To construct a plasmid (p*PAH1*) overexpressing *PAH1*, the DNA fragment containing own promotor and open reading frame of *PAH1* was amplified by PCR and cloned into pRS426 (2μ, *URA3*). The plasmid for expression of Opi1-GFP was constructed as follows. The BamHI-BglII fragment containing GFP(S56T) coding sequence from pFA6a-GFP(S65T)-kanMX6 was inserted into the BamHI site of pRS416 (CEN, *URA3*) to obtain pRS416-GFP. Subsequently, the EcoRI-BamHI fragment containing own promoter and open reading frame (without stop codon) of *OPI1* was amplified by PCR and cloned into pRS416-GFP to obtain p*OPI1*-GFP. To construct a plasmid expressing Kar2-SS-mRFP-HDEL, the DNA fragment containing Kar2 signal-peptide sequence (1-135)-mRFP-HDEL was amplified by PCR and cloned into pRS415 (CEN, *LEU2*) with GPD1 promotor. The plasmid for Yip3 overexpression was constructed by cloning the BamHI-HindIII fragment containing open reading frame of *YIP3* into pRS426 (2μ, *URA3*) with *GPD1* promoter.

### Culture conditions

Strains were grown either in rich YP medium (1% yeast extract, 2% peptone) supplemented with 0.2% adenine and containing 2% glucose (YPD) as carbon source or in synthetic dextrose (SD) minimal medium (0.15% yeast nitrogen base, 0.5% ammonium sulfate, 2% glucose) supplemented with the appropriate amino acids and bases as nutritional requirements or with 0.1% 5-fluoroorotic acid (5-FOA). To test the temperature sensitivity of strains, fivefold serial dilutions of cells were made in sterile water, spotted onto SD plates and incubated for 5 days (Kajiwara et al., 2012).

### Maturation of cargo proteins and protein levels of Sar1

Accumulation of immature CPY or Gas1 was analyzed by SDS-PAGE followed by immunoblotting (Ikeda et al., 2016). Blots were probed with rabbit polyclonal antibodies against CPY and Gas1, and detected by chemiluminescence using a peroxidase-conjugated affinity-purified goat anti-rabbit IgG antibody. Pulse-chase analysis for CPY maturation was carried out exactly as described (Kajiwara et al., 2014). The relative protein levels of Sar1 were analyzed by SDS-PAGE and immunoblotting using rabbit polyclonal antibodies against Sar1 and Wbp1 (a loading control).

### Fluorescence microscopy

For ERES visualization, cells expressing Sec13p-Venus were grown at 25°C in SD medium, shifted to 32 °C for 60 min and observed by conventional fluorescence microscopy. Cells were chosen randomly and ERESs were counted manually. For visualization of Opi1-GFP localization, cells were transformed with p*OPI1*-GFP. The cells were grown at 25°C in SD medium and imaged by fluorescence microscopy. Perinuclear ER/nucleoplasm intensity ratio was analyzed by Image J. At least 100 cells were evaluated for each sample per experiment. For ER membrane association of Sar1, cells expressing Sar1-AcGFP with Sar1-AcGFP (pRS316) plasmid (Yorimitsu and Sato, 2012) and an ER marker with Kar2-SS-mRFP-HDEL plasmid were grown at 25°C in SD medium, shifted to 32 °C for 60 min and imaged by fluorescence microscopy. For Sec16 localization, Sec16-GFP was expressed from the endogenous locus on chromosome, and Sec16-AcGFP was expressed from its own promoter on a pRS314-derived plasmid pTYY42-Sec16-AcGFP (CEN, *TRP1*) (Yorimitsu and Sato, 2012). ER membrane/cytosol intensity ratio was analyzed by Image J.

### DNA microarray analysis

Microarray analysis was performed as described previously (Wu et al., 2006), using the Gene Chip Yeast Genome 2.0 Array (Affymetrix). Total RNA was extracted from the yeast cells grown at 25°C in YPD medium and mRNA was purified using the Oligotex TM-dT30 mRNA Purification Kit (Takara Bio Inc., Japan). Biotinylated cRNA was prepared from 500 ng of mRNA according to the standard Affymetrix protocol, and 5 μg of cRNA was hybridized for 16 h at 45°C on the GeneChip Yeast Genome 2.0 Array. GeneChips were washed and stained using the Hybridization, Wash, and Stain Kit (Affymetrix). Data were analyzed with Operating Software (GCOS) v1.4, using the Affymetrix default analysis settings and global scaling as the normalization method. The trimmed mean target intensity of each array was arbitrarily set to 500. Microarray data can be retrieved from Gene Expression Omnibus (GEO) under the accession code GSE168638. DNA microarray data were obtained for three independent culture experiments from *sec12-4* and *sec12-4 ino2*Δ *ino4*Δ cells.

### MS analysis of glycerophospholipids

Mass spectrometer (MS) analysis of glycerophospholipids was carried out exactly as described (Melero et al., 2018). Cells were grown at 32°C in SD medium and the lipids were extracted with the extraction solvent (ethanol, water, diethyl ether, pyridine, and 4.2 N ammonium hydroxide [15:15:5:1:0.018, vol/vol]) with internal standard mix. Glycerophospholipids were analyzed by using a Triple Stage Quadrupole (TSQ) Vantage Mass Spectrometer (Thermo Scientific) equipped with a robotic nanoflow ion source Nanomate HD (Advion Biosciences, Ithaca, NY). The abundance of glycerophospholipids was calculated by their signal intensities relative to the internal standards and standard curves constructed using the internal standard lipids. The data presented are not corrected for +2 isotopes as the data are presented as relative to wild type and vast majority of the isotope correction would be cancelled out.

### TLC analysis of phospholipids

Phospholipids extracted from cells were spotted onto thin-layer chromatography (TLC) plates that were pretreated with 2.3% (w/v) boric acid in 100% ethanol for at least 15 min and dried in 100°C oven. The loaded TLC plates were developed with a solvent system containing chloroform/ethanol/water/triethylamine (30:35:7:35). The plate was air-dried and the lipids were detected by ultraviolet light after spraying with 0.05% primulin in acetone/water (8/2, v/v).

### Statistical analysis

Statistical analysis was performed using Student’s t-Test or Tukey-Kramer multiple comparison test (ns not significant; *, p < 0.05; **, p < 0.01 and ***, p < 0.001). The mean ± s. d. for three independent experiments is shown.

## DATA AVAILABILITY

DNA microarray data for *sec12-4* and *sec12-4 ino2Δ ino4Δ* have been deposited under GEO Accession Number GSE168638 in NCBI and provided as a supplemental dataset (Dataset EV3). Lipidomics data related to this study are provided as supplementary datasets (Datasets EV2 and EV4).

## ACKNOWLEDGMENTS

We thank R Schekman for yeast *sec* mutant strains, the National Bio-Resource Project/Yeast Genetic Resource Center (NBRP/YGRC) of Japan for yeast strains with gTOW plasmids, and K. Sato for Sar1-AcGFP, Sar1D32G-AcGFP (pRS316), and pTYY42-Sec16-AcGFP (pRS314) plasmids. This work was funded by the Grants-in-Aid for Scientific Research from Japan Society for the Promotion of Science, Japan [JP19H02922, JP21K19088 to K.F.], by the FEDER/Ministerio de Ciencia, Innovación y Universidades-Agencia Estatal de Investigación/BFU2017-89700-P and PID2020-119505GB-I00 to M.M., by the Junta de Andalucía, Spain PY20_01240 to M.M., and by the “VI Own Research Plan” of the University of Seville VIPPIT-2020-I.5 to M.M., and by the SNSF (grants 51NF-40-185898 and 310030_184949) and the Leducq Foundation to H.R..

## AUTHOR CONTRIBUTIONS

M. N., H. N., M. K., R. I., M. I., K. H., K. E., T. K., A. I., and P. S. constructed strains and plasmids, and performed biochemical studies and fluorescent microscopy experiments and data analysis.

J. M-L., A. A-R., S. S-B. and A. P-L. performed pulse-chase experiments with help from M. M..

M. K., A.I., Y. Y., and M. K. performed microarray experiments with help from H. I..

I. R. and H. R. performed MS analysis for lipids.

K. F designed the experiments and wrote the manuscript with input from all other authors.

## EXPANDED VIEW FIGURE LEGENDS

**Appendix Figure S1.**
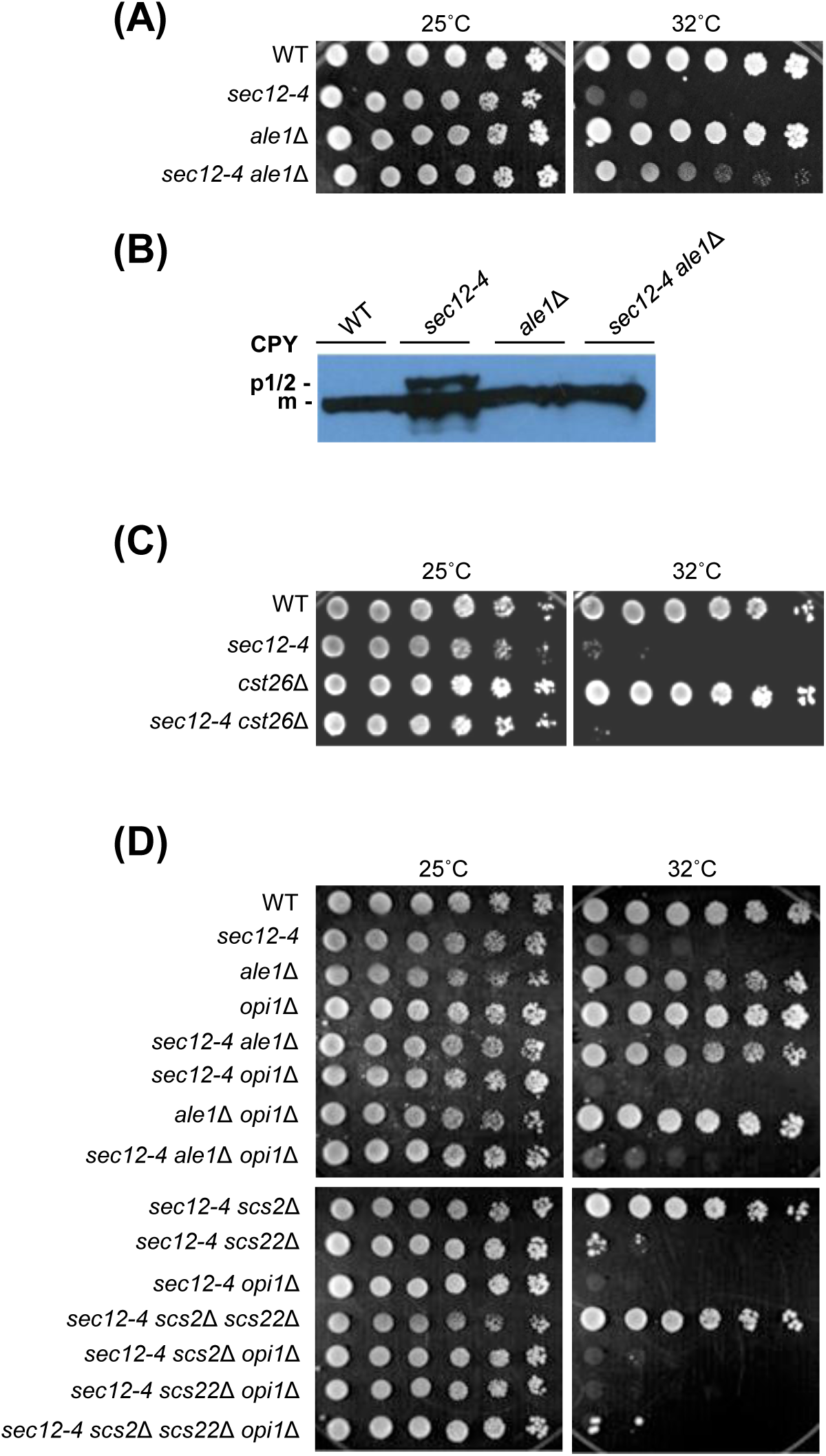
Deletion of *ALE1*/*SLC4* or *SCS2*/*SCS22* but not *CST26* rescues *sec12-4* mutant phenotypes and Opi1 is required for the rescue, Related to Figure 1 and 2. (A-D) Spot assay (A, C and D) and western blot analysis (B) were performed as described in Figure 1.

**Appendix Figure S2.**
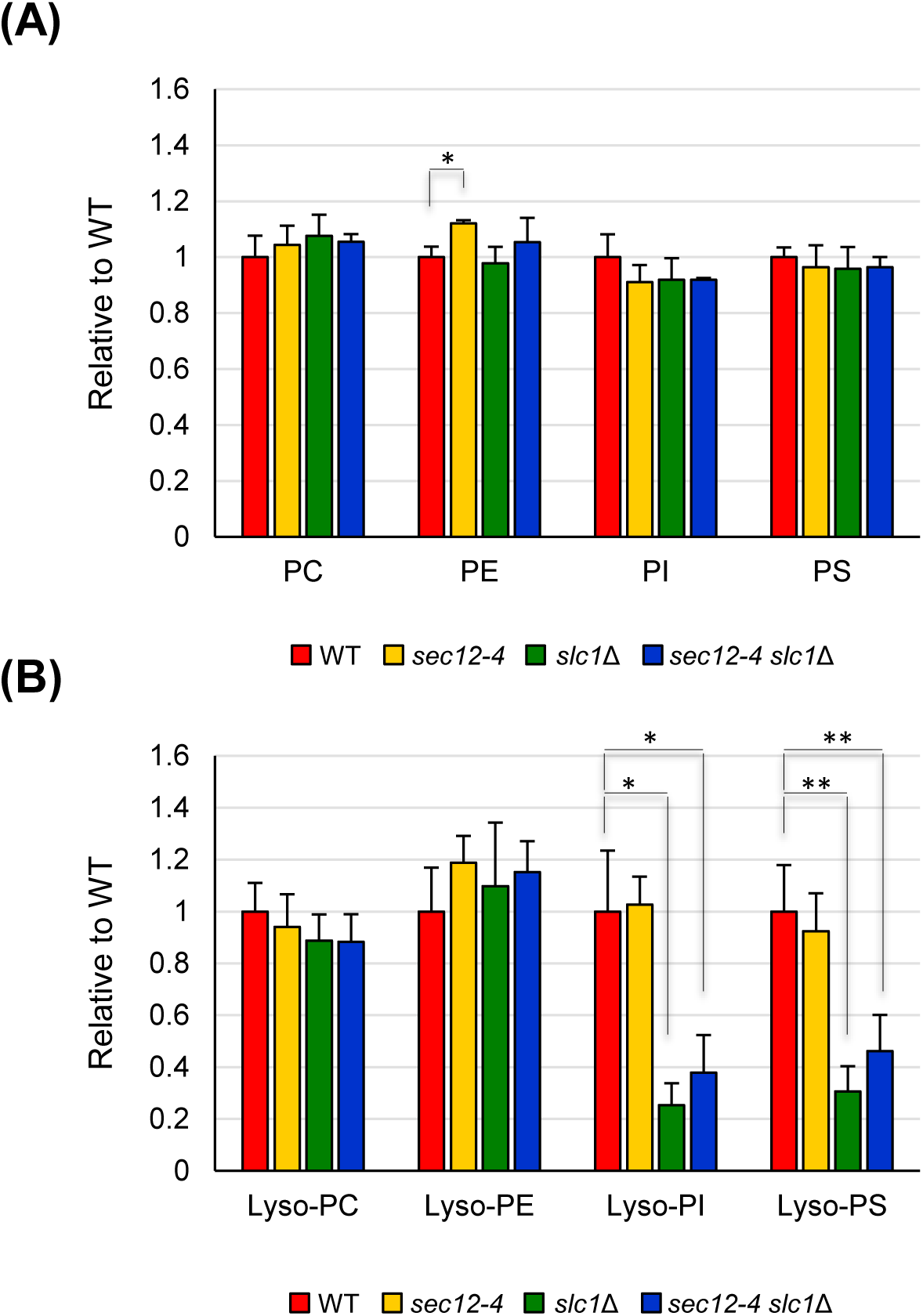
Lipid profiles of main glycerophospholipids species (upper histogram, A) and their lysolipid counterparts (lower histogram, B), Related to Figure 1. Cells were grown at 32°C and glycerophospholipids were extracted and analyzed by mass spectrometer. Lipid species were identified according to their m/z, and their abundance was calculated by their signal intensities relative to internal standards. The relative amounts of lipids from wild type and mutant cells were calculated and wild type was set at 1. The results represent the averages of three independent experiments, and standard deviations are included. Student’s t test: **p ≤ 0.01, *p ≤ 0.05.

**Appendix Figure S3.**
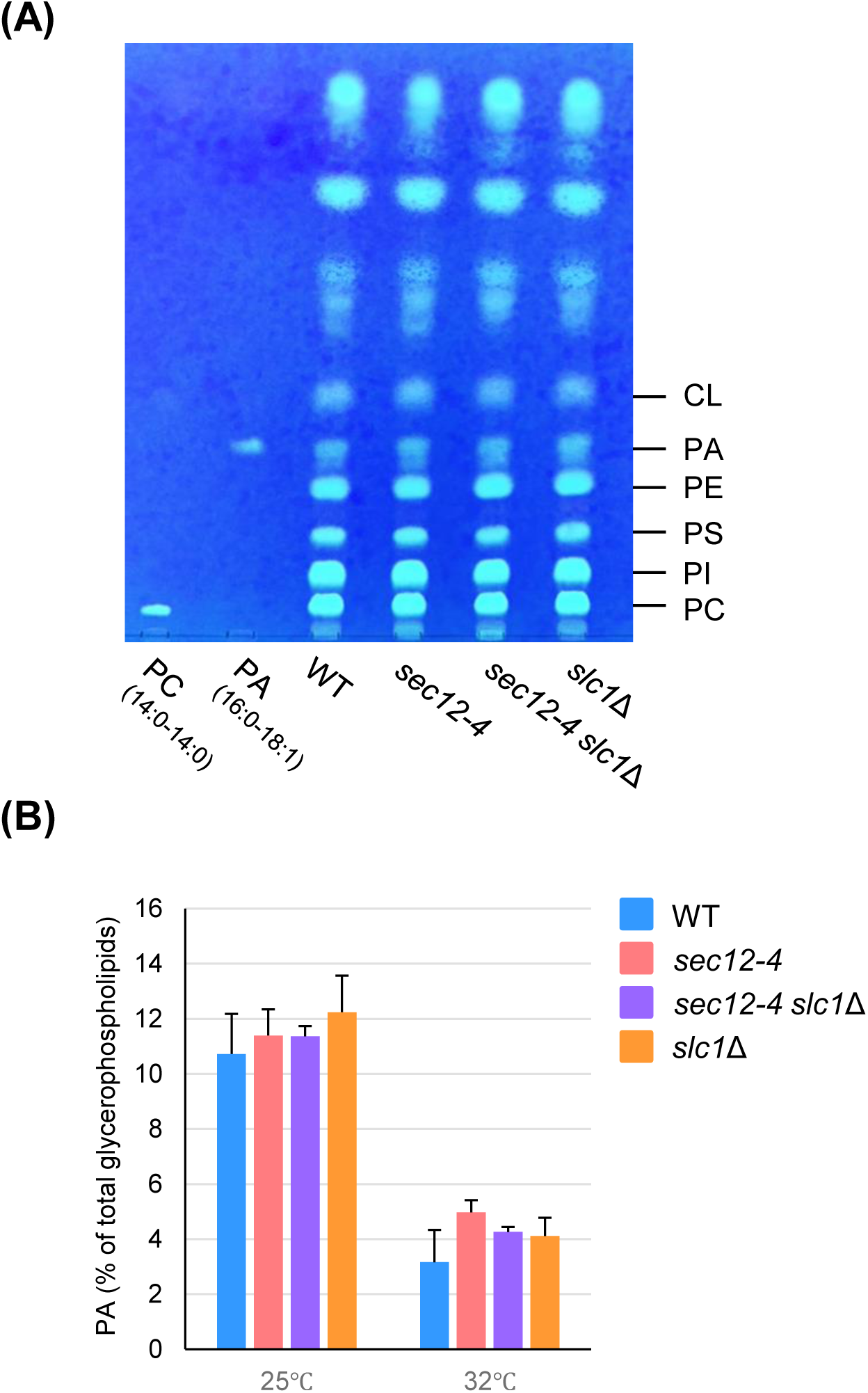
Loss of Slc1 does not affect the total level of phosphatidic acids, Related to Figure 1. Phospholipids extracted from cells grown at 25°C (A, B) or 32°C (B) were separated by TLC and visualized by spraying with a solution of primulin followed by illumination with UV light. Phosphatidic acids (PA) were identified by comparison with a purified reference standard, PA (16:0-18:1). Percentages of PA to total phospholipids were measured and the results represent the averages of three independent experiments, and standard deviations are included.

**Appendix Figure S4.**
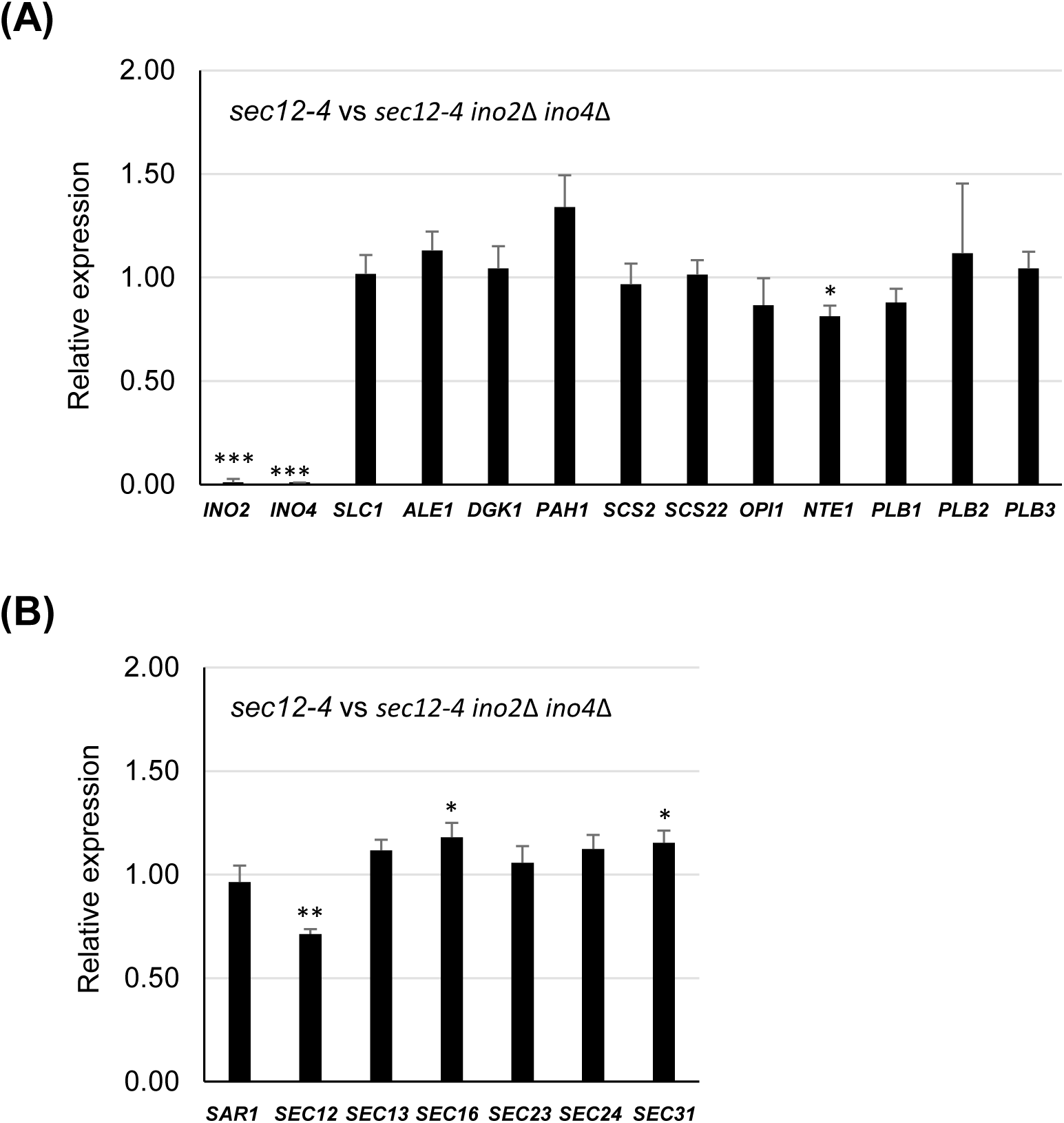
Relative expression levels of genes involved in glycerophospholipid metabolism and COPII vesicle formation, Related to Figure 4. The relative expression levels of genes involved in glycerophospholipid metabolism (A) and COPII vesicle formation (B) were determined by microarray analysis in RNA samples obtained from *sec12-4* and *sec12-4 ino2*Δ *ino4*Δ cells. Mean values and standard deviations were obtained from three independent experiments. Student’s t test: ***p ≤ 0.001, **p ≤ 0.01, *p ≤ 0.05.

**Appendix Figure S5.**
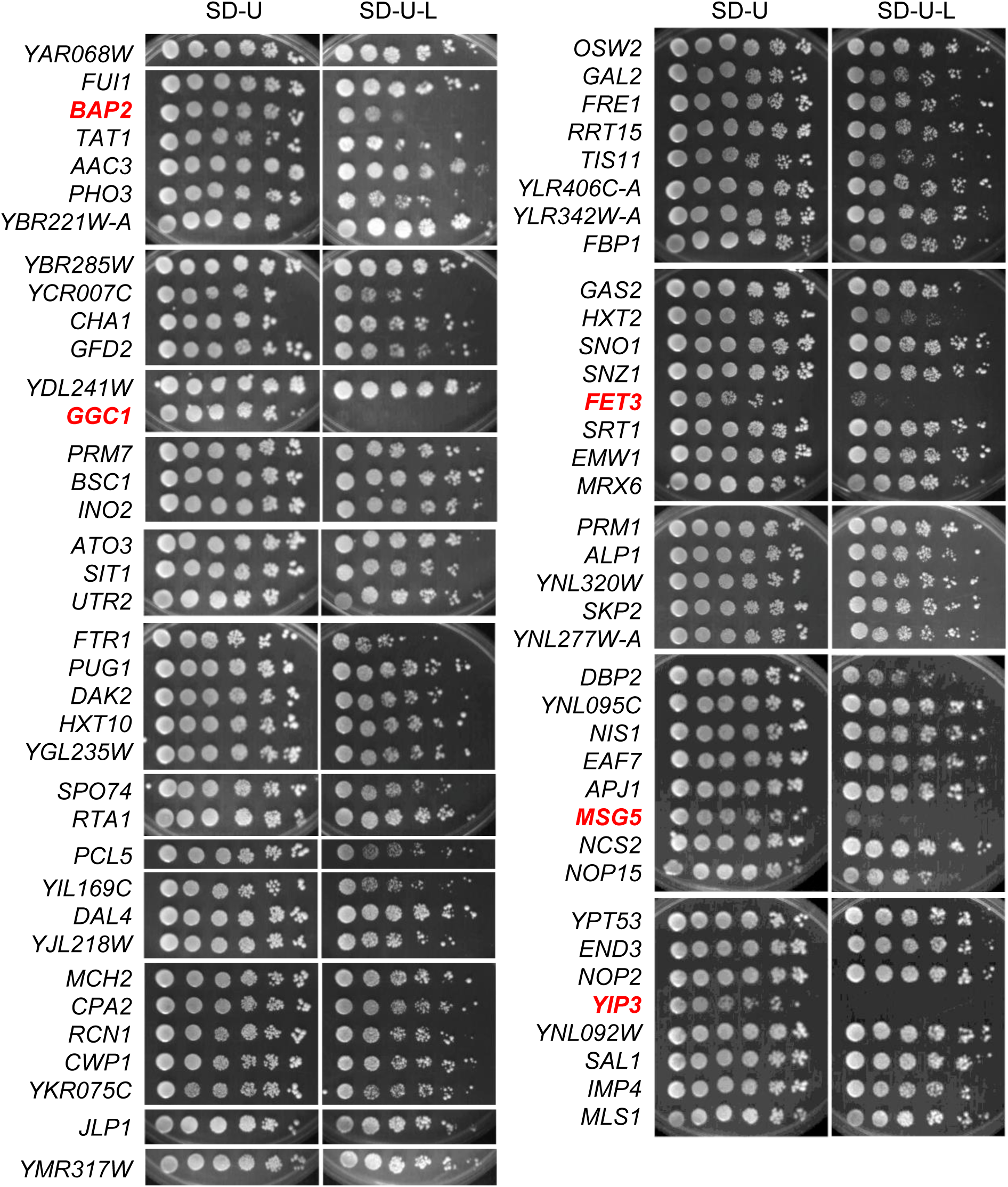
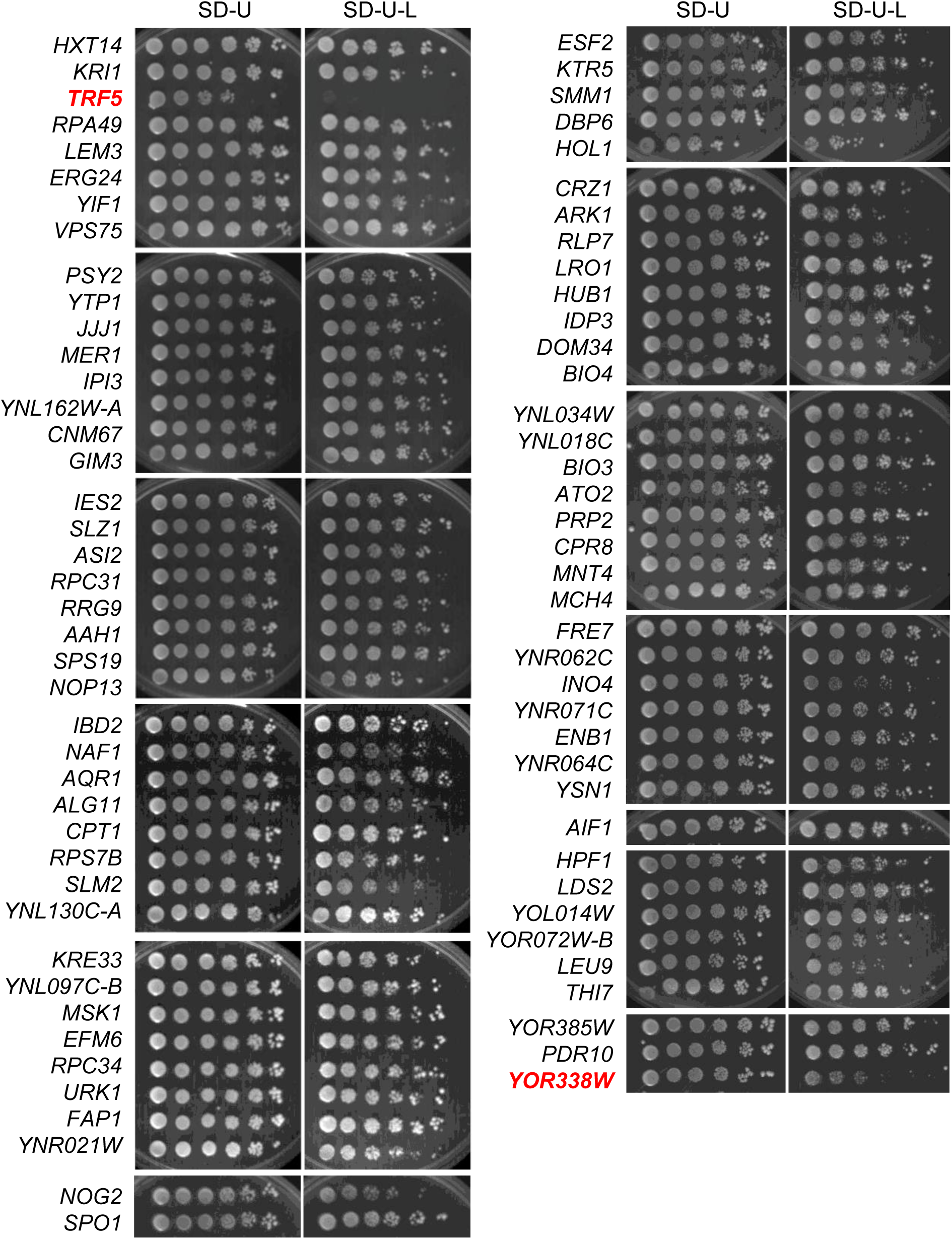
Identification of genes that cause dosage sensitivity as negative regulators of COPII vesicle formation, Related to Figure 4. A screening to identify dosage sensitive genes was performed using the genetic tug-of-war (gTOW) method. Cells bearing gTOW plasmids were grown at 25°C in SD medium lacking uracil, the five-fold serial dilutions were spotted onto SD plates without uracil (SD-U) or both uracil and leucine (SD-U-L) and were incubated at 25°C for 5 days. 7 genes that cause a growth defect when overexpressed in SD-U-L are shown in red.

**Appendix Figure S6.**
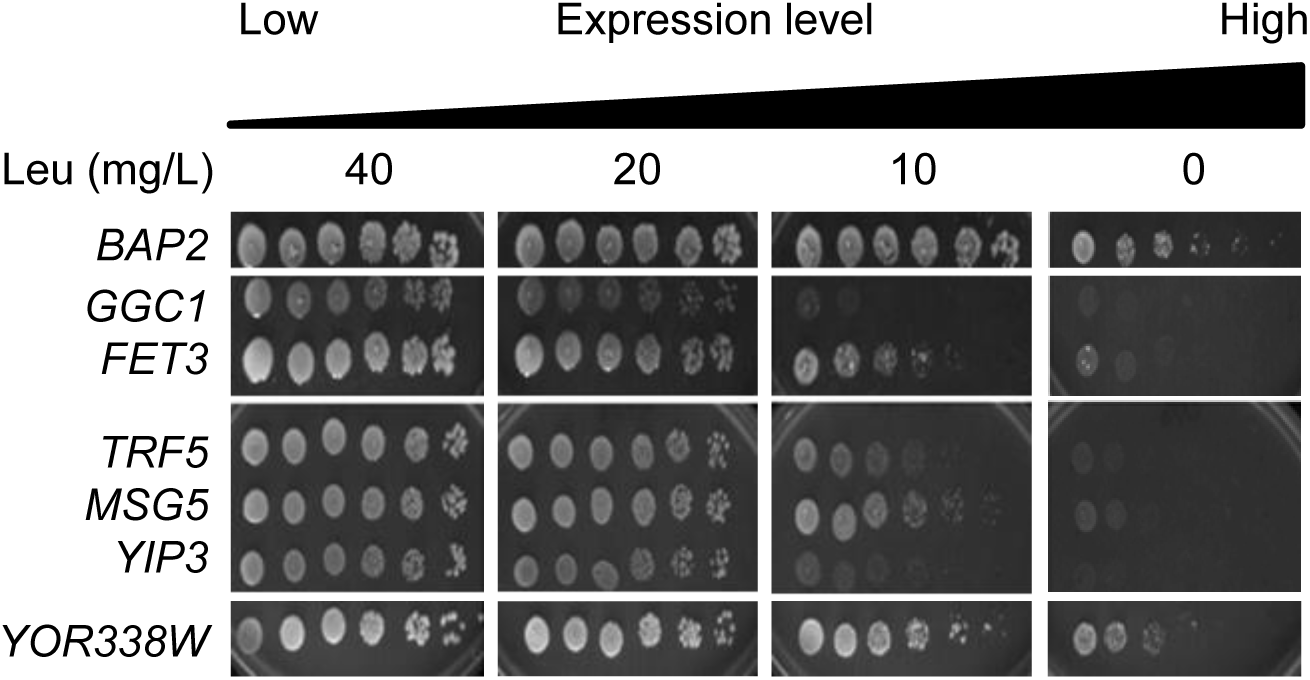
7 Candidate genes cause dosage sensitivity depending on leucine concentration in medium, Related to Figure 4. Cells bearing gTOW plasmids were grown at 25°C in SD medium lacking uracil, the five-fold serial dilutions were spotted onto SD plates without uracil (SD-U) and the indicated concentrations of leucine and were incubated at 25°C for 5 days.

**Appendix Figure S7.**
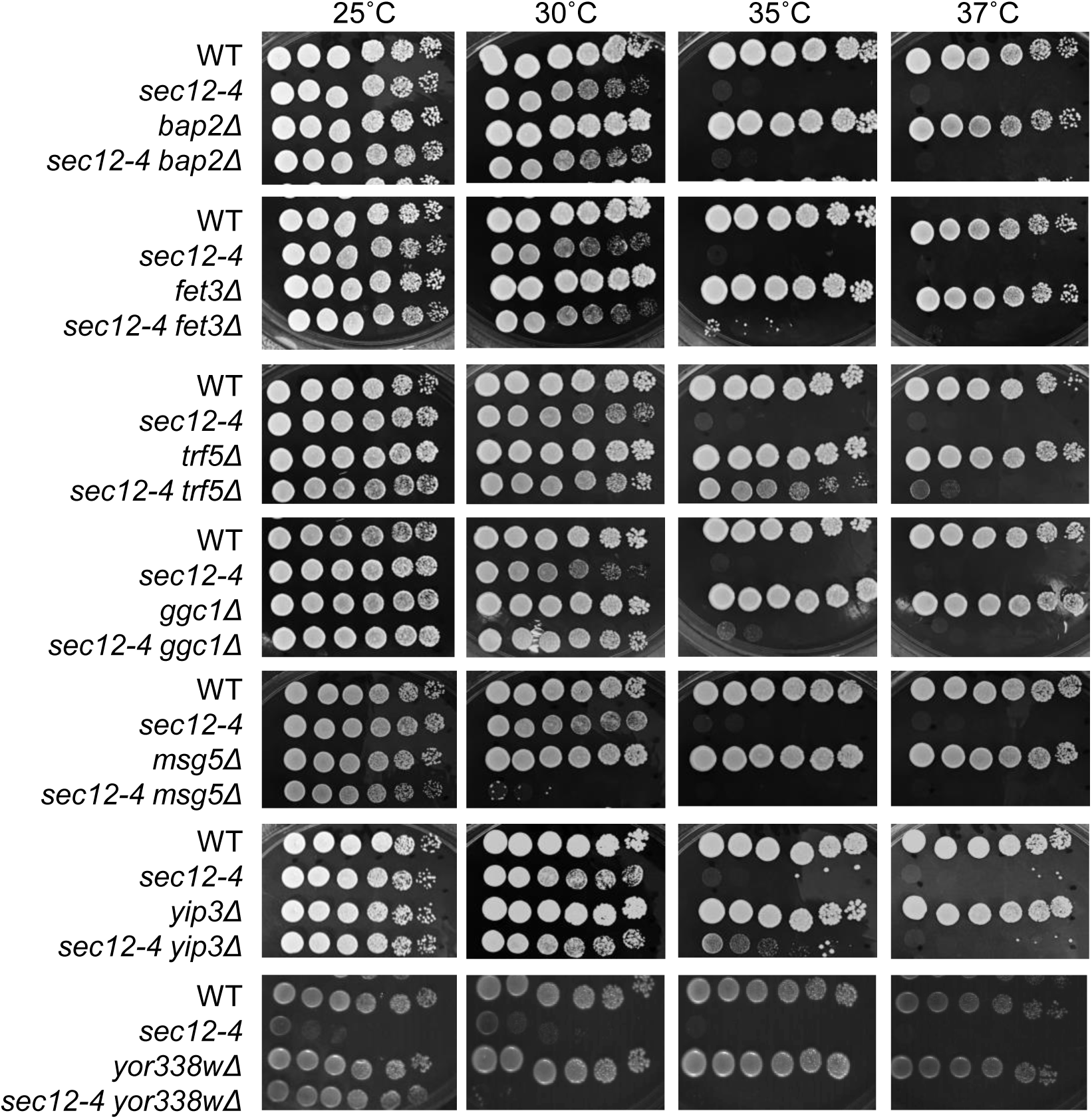
Deletion of *TRF5* or *YIP3* rescues TS growth defect of *sec12-4* mutant, Related to Figure 4. Spot assay was performed as described in Figure 1.

**Appendix Figure S8.**
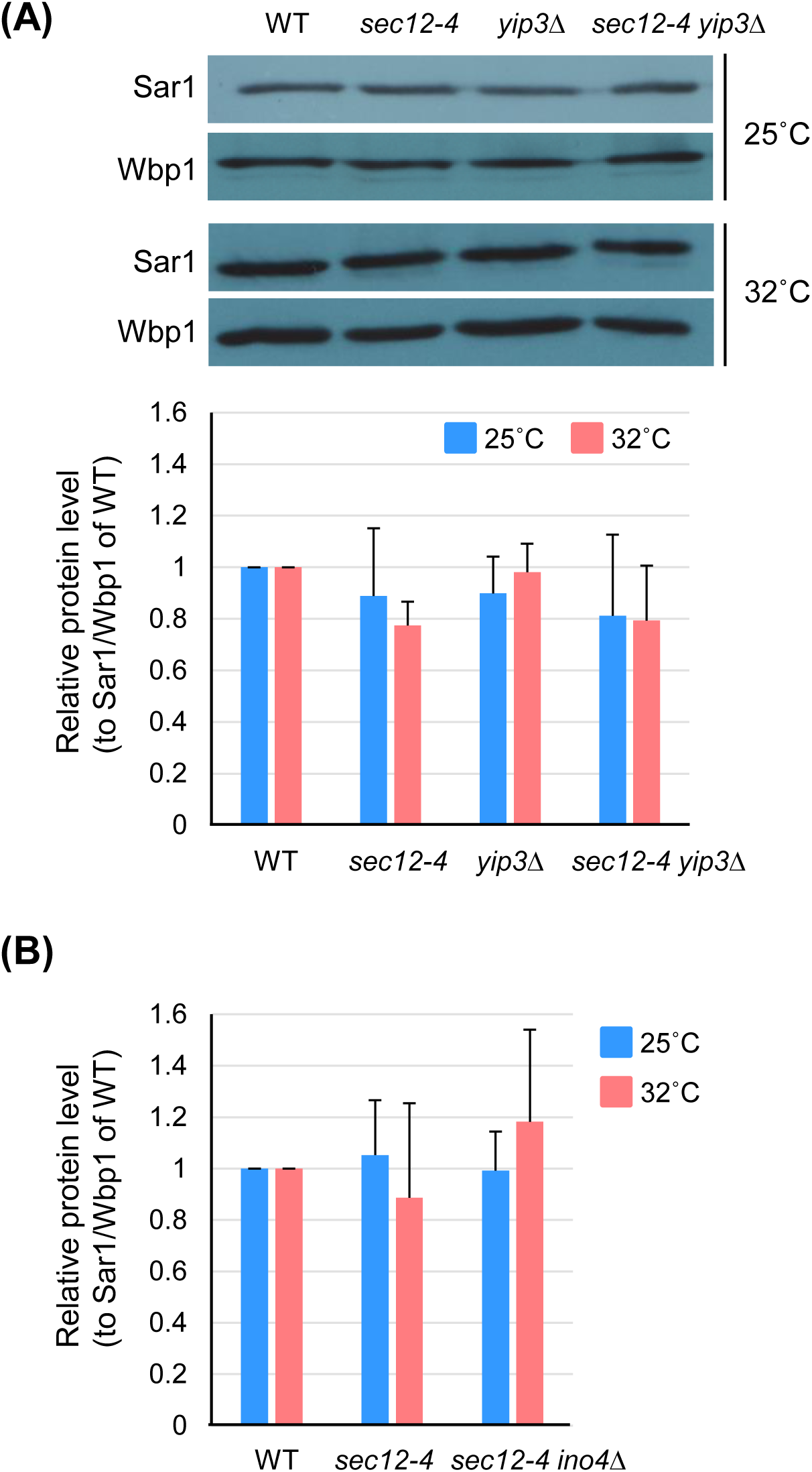
Protein levels of Sar1 were unaffected by *YIP3 or INO4* deletion, Related to Figure 5. The relative steady-state protein levels of Sar1 were determined by western blot analysis. Cells grown at 25°C were shifted to 32 °C for 90 min, and protein extracts were analyzed. Wbp1 was used as a loading control. Mean values and standard deviations of the relative amount to the ratio of Sar1/Wbp1 in wild-type cells were obtained from three independent experiments.

**Appendix Figure S9.**
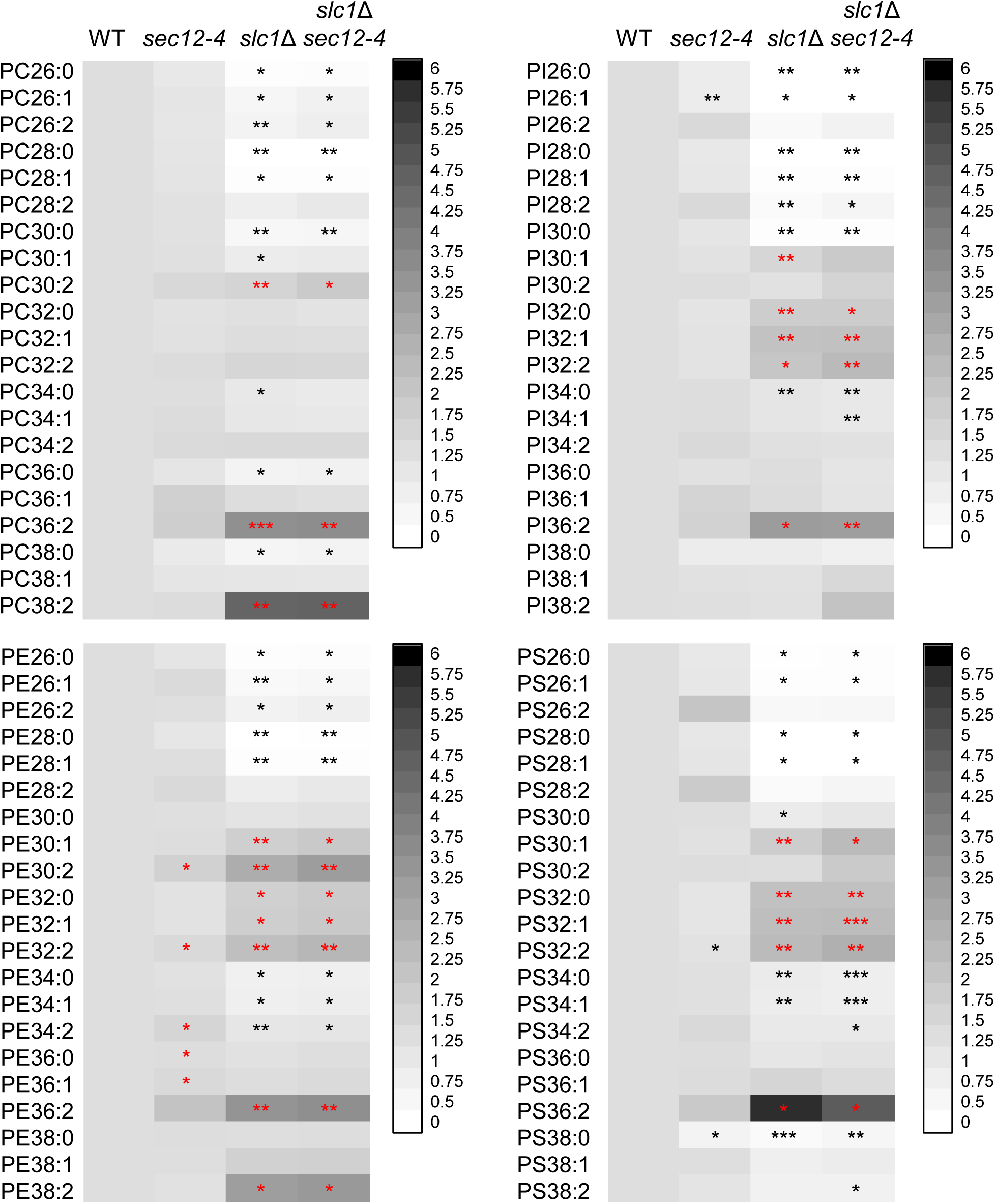
Fatty acyl-based analysis of phospholipids in *sec12-4*, *slc1*Δ and *sec12-4 slc1*Δ mutants, Related to Figure 1. Analysis of phospholipids was performed as described in Figure S1. The quantities of lipids were expressed as signal intensities relative to wild type levels and data are presented as averages of three independent experiments and in heat map visualizations. Statistical significance between wild type and *slc1*Δ or *sec12-4* and *sec12-4 slc1*Δ cells was determined. Student’s t test: ***p ≤ 0.001, **p ≤ 0.01, *p ≤ 0.05. Significant decreases are indicated by black asterisks, and significant increases by red asterisks.

### Supplementary datasets

**Dataset EV1.** Strains of *Saccharomyces cerevisiae* used for this study.

**Dataset EV2.** Average and SD values for the lipid composition, related to Figure S2.

**Dataset EV3.** Average and SD values for the transcriptomic profiles of *sec12-4* and *sec12-4 ino2*Δ *ino4*Δ mutant cells, related to Figure 4A.

**Dataset EV4.** Average and SD values for the lipid composition, related to Figure S9.

